# Indoleamine 2,3-dioxygenase upregulates PD-1 expression on ovarian tumor infiltrating CD8^+^ T cells via kynurenine activation of the aryl hydrocarbon receptor

**DOI:** 10.1101/2021.02.18.431473

**Authors:** Adaobi Amobi-McCloud, Ravikumar Muthuswamy, Sebastiano Battaglia, Han Yu, Tao Liu, Jianmin Wang, Vasanta Putluri, Prashant K Singh, Feng Qian, Ruea-Yea Huang, Nagireddy Putluri, Takemasa Tsuji, Amit A Lugade, Song Liu, Kunle Odunsi

**Affiliations:** Department of Immunology, Roswell Park Comprehensive Cancer Center, Buffalo, NY; Center for Immunotherapy, Roswell Park Comprehensive Cancer Center, Buffalo, NY; Advanced Technology Cores, Baylor College of Medicine, Houston, TX; Department of Molecular and Cell Biology, Baylor College of Medicine, Houston, TX; Department of Biostatistics & Bioinformatics, Roswell Park Comprehensive Cancer Center, Buffalo, NY; Center for Personalized Medicine, Roswell Park Comprehensive Cancer Center, Buffalo, NY; Department of Gynecologic Oncology, Roswell Park Comprehensive Cancer Center, Buffalo, NY

## Abstract

The immunoregulatory enzyme, indoleamine 2,3-dioxygenase (IDO1) and the PD-1/PD-L1 axis are potent mechanisms that impede effective anti-tumor immunity in ovarian cancer. However, whether the IDO pathway regulates PD-1 expression in T cells is currently unknown. Here we show that tumoral IDO1 expression led to profound changes in tryptophan, nicotinate/nicotinamide, and purine metabolic pathways in the ovarian tumor microenvironment, and to an increased frequency of PD-1^+^CD8^+^ tumor infiltrating T cells. We determined that activation of the aryl hydrocarbon receptor (AHR) by kynurenine induced PD-1 expression, and this effect was significantly abrogated by the AHR antagonist CH223191. Mechanistically, kynurenine alters chromatin accessibility in regulatory regions of T cell inhibitory receptors, allowing AHR to bind to consensus XRE motifs in the promoter region of PD-1. These results enable the design of strategies to target the IDO1 and AHR pathways for enhancing anti-tumor immunity in ovarian cancer.

## Introduction

Epithelial ovarian cancer (EOC) is the most lethal gynecologic malignancy in the U.S. (1). Despite initial response to frontline treatments (i.e. surgery and platinum-based chemotherapy), the majority of patients relapse and ultimately die from their disease within five years (2, 3). Although several studies have demonstrated a positive correlation between EOC prognosis and magnitude of tumor-infiltrating effector T lymphocytes (TIL) (4–6), the clinical benefit of TIL-promoting immunotherapies – such as immune checkpoint inhibitors (ICI), vaccines, and adoptive cell therapy – is limited by the presence of multiple adaptive and synergistic immune tolerance mechanisms within the ovarian tumor microenvironment (TME).

Among several immunosuppressive mechanisms, indoleamine 2,3-dioxygenase (IDO1) has emerged as a key targetable pathway impacting the anti-tumor function of TILs. IDO1 is a heme enzyme which catabolizes the first and rate-limiting step of tryptophan (Trp) catabolism along the kynurenine pathway (KP) to generate active immunosuppressive metabolites. Depletion of tryptophan leads to arrest of T cell proliferation (7) by inhibiting the mechanistic target of rapamycin complex 1 (mTORC1) (8), and inducing a stress response via activation of the general control nondepressible-2 (GCN2) kinase (9). In addition, kynurenine (Kyn) promotes the differentiation of CD4^+^ T cells into immunosuppressive regulatory T (Treg) cells via activation of the aryl hydrocarbon receptor (AHR) (10, 11). In EOC patients, elevated IDO1 expression correlated with a lower Trp:Kyn ratio in the ovarian tumor microenvironment (12), reduced CD8^+^ TIL frequency (13), poor prognosis (14, 15), and suppression of T cell responses (16). The vital role of targeting IDO1 for effective immunotherapeutic control of established tumors was observed in pre-clinical models by the synergistic effect of IDO1 inhibition and immune checkpoint inhibitors to mediate the rejection of poorly immunogenic tumors, indicating that IDO1 may be a major resistance mechanism (17).

Although these observations support therapeutic targeting of the IDO1 pathway, EOC patients treated with epacadostat, an IDO1 inhibitor, did not exhibit objective responses with a median progression-free survival (PFS) of 3.75 months versus 5.56 months for the control group receiving tamoxifen (18). Moreover, a subsequent randomized phase III clinical trial in patients with unresectable metastatic melanoma (19) failed to demonstrate improvement in clinical responses when epacadostat was added to pembrolizumab (20–22). These findings suggest that a gap still exists in understanding the full biological consequences of IDO1 enzyme activity in the TME.

Since high IDO1 enzyme activity (12) occurs concomitantly with elevated PD-1 expression on antigen-specific CD8^+^ T cells as a marker of exhaustion and dysfunction (23), we reasoned that IDO1 may play a role in regulating the expression of PD-1 and other inhibitory receptors in EOC. As the IDO1 metabolite Kyn is an endogenous ligand of AHR transcription factor (24), we investigated a possible role for AHR as the mechanism by which IDO1 facilitates TIL dysfunction associated with inhibitory checkpoint receptor upregulation. In this study, we observed profound metabolic and immunoregulatory changes in the ovarian TME based on IDO1 expression, and importantly, induction of inhibitory receptors on CD8^+^ TILs via Kyn-mediated AHR signaling. These data implicate a novel role for Kyn in regulating the exhausted phenotype of CD8^+^ T cells.

## Results

### IDO1 reduces the prognostic benefit of TILs in human EOC and impacts overall survival

We evaluated the clinical outcome of 265 patients with high-grade serous ovarian cancers available in TCGA stratified by TIL expression and 44 genes (Supplemental Table 1) related to tryptophan catabolism and AHR signaling. TCGA EOC patient cohorts stratified into four distinct populations (TIL^High^/IDO^Low^, TIL^Low^/IDO^Low^, TIL^Low^/IDO^High^, and TIL^High^/IDO^High^) (Figure 1A). TIL^High^/IDO^Low^ patients had a significantly improved disease-free survival (DFS) and overall survival (OS) compared with the other groups (Figure 1B). Additionally, elevated IDO1 and AHR pathway expression negated the beneficial impact of increased TIL signature (TIL^High^/IDO^High^ patients), further highlighting a critical role for this pathway. These data suggest that the relationship between IDO1 expression and TIL infiltration is critical in shaping EOC patient outcomes.

**Figure 1:**
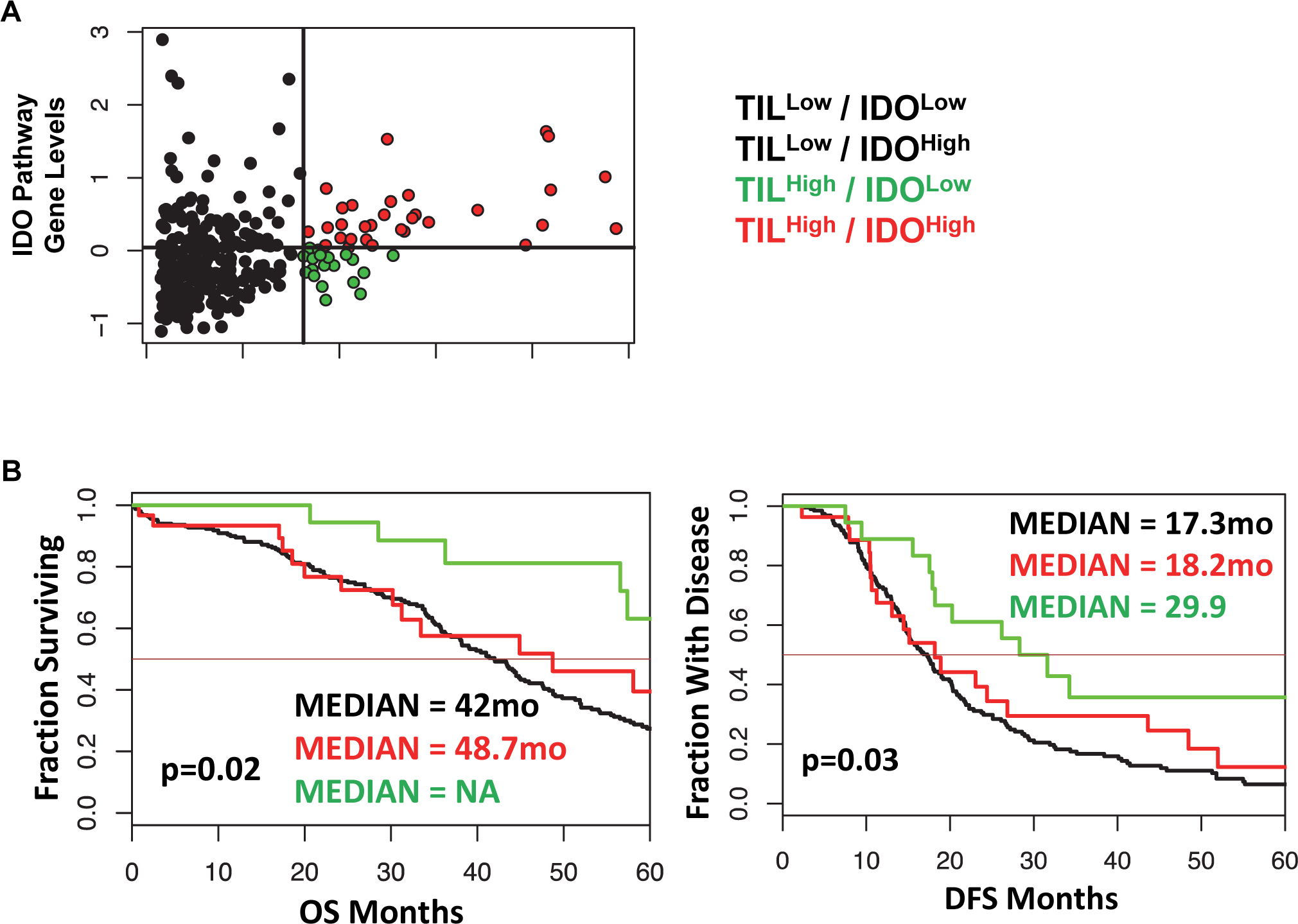
IDO1 reduces the prognostic benefit of tumor infiltrating CD8^+^ T cells in human ovarian cancer and reduces overall survival in a murine model of ovarian cancer. **(A)** Scatterplot and **(B)** Kaplan-Meier curves of 4 distinct populations comprised of 265 high grade serous ovarian cancer patients from The Cancer Genome Atlas (TCGA) data set. RNA-seq data was analyzed in the context of 44 genes from the tryptophan metabolism and AHR signaling pathways, and CD3E, CD8A, IL-2, and Granzyme B. Confidence intervals for the stratified population of patients include OS: black line median 42.0[38.0, 46], red line median 48.7[31.2, NA] and green line median NA[57.4, NA] *p*=0.02; DFS: black line median 17.3[15.1, 19.9], red line median 18.2[13.0, 48.5], and green line median 29.9[18.2, NA] *p*=0.03). **(C)** 6- to 8-week old WT C57BL/6 mice challenged intraperitoneally (i.p.) with 1×10^7^ IE9mp1-EV (n=10) or IE9mp1-mIDO1 (n=12) tumor cells. Tumor progression was quantified by measuring the abdominal circumference of tumor-bearing mice. **(D)** Kaplan-Meier curves of the survival analysis of IE9mp1-EV (n=14) and IE9mp1-mIDO1 (n=15) tumor-bearing WT C57BL/6 mice. ∗p < 0.05, ∗∗∗p < 0.001, by the Log-rank (Mantel-Cox) test (**B** and **D**), or Student’s t test **(C)**.

To delineate the mechanisms by which IDO1 mediates immune suppression, we generated a stable IDO1-expressing EOC cell line by retroviral transduction (Supplemental Figure 1A) of an aggressive ID8 variant, IE9mp1 (25, 26). IE9mp1-mIDO1 tumor cells expressed the murine IDO1 (mIDO1) gene (Supplemental Figure 1B) and the gene product demonstrated functional enzyme activity, as measured by elevated Kyn production compared to empty vector (IE9mp1-EV) controls (Supplemental Figure 1C). The addition of the mIDO1 gene did not alter *in vitro* cell viability compared with EV control (Supplemental Figure 1D). Consistent with the TCGA EOC data, syngeneic wild-type (WT) C57BL/6 mice challenged with IE9mp1-mIDO1 displayed earlier onset of tumor burden (Figure 1C) and a significant decrease in overall survival compared with tumors that lack IDO1 expression (Figure 1D).

### Expression of IDO1 profoundly alters the metabolic profile of ovarian tumors

Dynamic changes in the metabolic profile of intraperitoneal IDO1-expressing ovarian tumors were evaluated by LC/MS measurement of Kyn and its downstream catabolites (27). Unsupervised principal component analysis (PCA) showed a differential metabolite signature of IDO1-expressing tumors (red triangles) that altered during tumor progression from day 28 to day 47 (Figure 2A). IDO1 expression at endpoint (Day 47) had the largest effect on the tryptophan, nicotinate/nicotinamide, and purine metabolism pathways (p<0.1; Global-ANCOVA) (Table 1). As expected, IDO1-expressing tumors demonstrated lower levels of tryptophan compared to EV tumors, alongside elevated expression of downstream Kyn metabolites (Figure 2B and Supplemental Figure 2A). The metabolite signature also revealed elevation in nicotinic acid, nicotinamide, and quinolinic acid (Figure 2C and Supplemental Figure 2A), consistent with enhanced *de-novo* nicotinamide generation via the Kyn pathway (28, 29). Nicotinate and nicotinamide metabolites (which include adenosine and thymine) were increased in the IE9mp1-mIDO1 tumors compared with IE9mp1-EV tumors at day 47 (Figure 2C, Supplemental Figures 2B and 2C). Altogether, the impact of tumoral IDO1 expression in ovarian cancer was not only confined to the kynurenine pathway, but also affected nicotinamide, purine and pyrimidine metabolic pathways, and the magnitude of change was influenced by the tumor burden.

**Table 1:** Global-ANCOVA analysis of IE9mp1-mIDO1 or IE9mp1-EV tumor from WT C57BL/6 mice (p<0.1). P-value indicates the significance of overall change for the metabolic pathway in IDO1-expressing tumors. Impact score measures the importance of a metabolite in the metabolic network.

| Pathway | Total<br>Cmpd | Hits | p-value | Impact |
| --- | --- | --- | --- | --- |
| ***Tryptophan metabolism*** | **79** | **5** | **0.003523** | **0.26445** |
| ***Nicotinate and nicotinamide***<br>***metabolism*** | **44** | **5** | **0.029416** | **0.1684** |
| ***Nitrogen metabolism*** | **39** | **4** | **0.04472** | **0.00067** |
| *Purine metabolism* | 92 | 12 | 0.063504 | 0.20329 |
| *Pyrimidine metabolism* | 60 | 5 | 0.43918 | 0.19526 |

**Figure 2:**
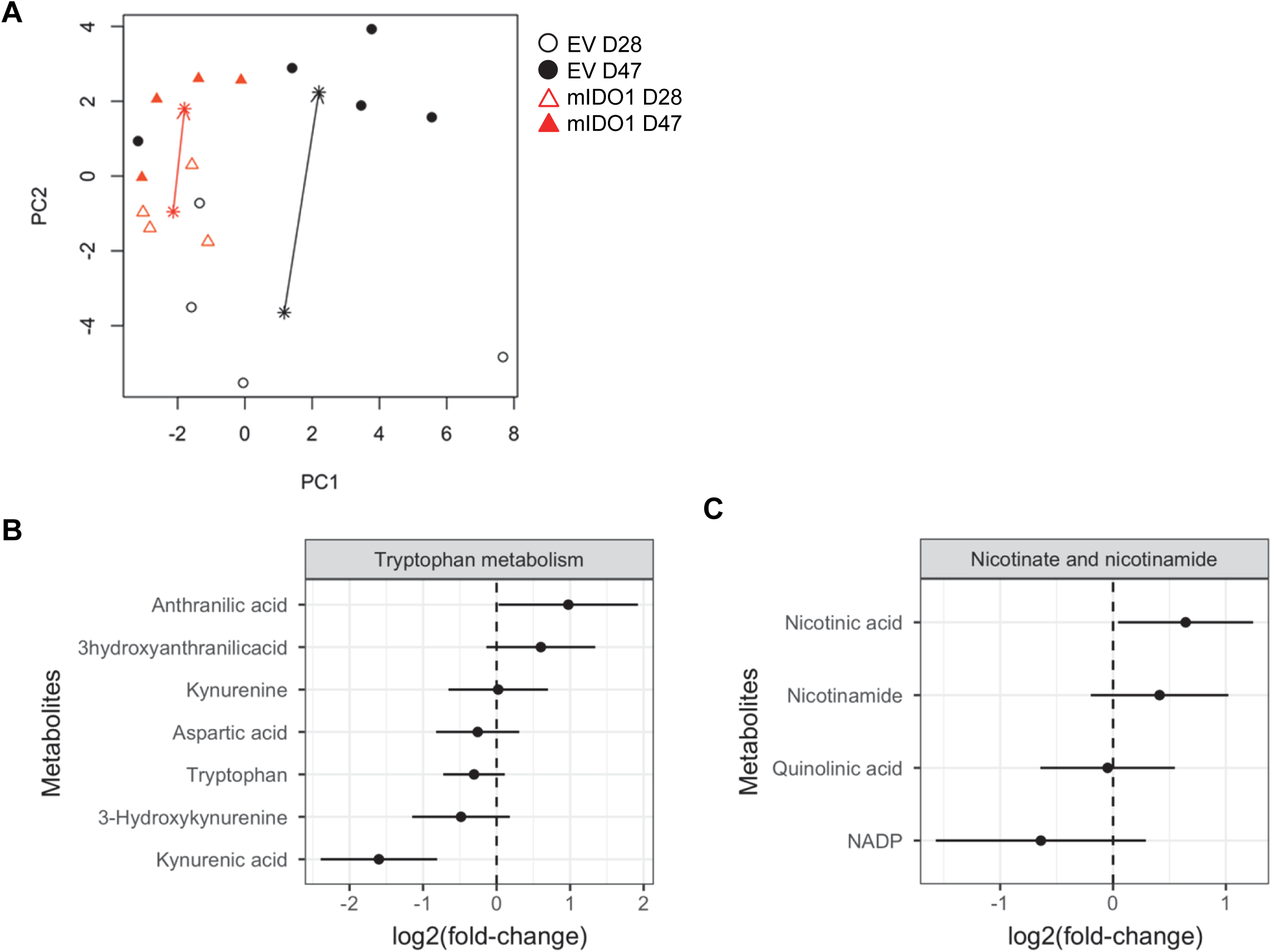
Tumoral IDO1 expression alters kynurenine, nicotinamide, purine and pyrimidine metabolic pathways. **(A)** Principal component analysis of metabolites measured in IE9mp1-mIDO1 (red triangles) and IE9mp1-EV (black circles) tumors from WT C57BL/6 mice at Day 28 (open) and Day 47 (solid). Asterisks are means of time-treatment group and arrows show the change in means from Day 28 to 47. **(B)** The 95% confidence interval of mean log-fold change in kynurenine, 3-hyrdroxyanthranilic acid, anthranilic acid, **(C)** quinolinic acid, nicotinic acid and nicotinamide metabolites levels in IE9mp1-mIDO1 versus IE9mp1-EV tumor tissue at Days 28 and 47 are shown.

### Host- and tumor-derived IDO1 expression drive an immunosuppressive cell profile in the ovarian TME

To delineate the relative contribution of host-versus tumor-derived IDO1 on the TME immune cell profile, IDO1-sufficient C57BL/6 (WT) mice and IDO1-knockout mice (IDOKO, C57BL/6 background) were challenged intraperitoneally with either IE9mp1-EV or IE9mp1-mIDO1. WT mice bearing IE9mp1-mIDO1 tumors exhibited decreased CD8^+^ TIL frequency at all time points of tumor growth compared to animals whose tumors lack IDO1 expression (Figure 3A, left panel). Genetic restriction of IDO1 expression to tumors (i.e. IDOKO mice) also resulted in diminished CD8^+^ TIL frequency (Figure 3A, right panel), but the complete absence of IDO1 expression in tumor and host increased CD8^+^ TIL frequency, indicating that both host- and tumor-derived IDO1 contribute to reduced TIL accumulation in the ovarian TME. Confocal microscopy of IE9mp1-mIDO1 and IE9mp1-EV tumors also confirmed that tumor-derived IDO1 inversely impacted CD8^+^ TIL frequency (Figure 3B).

**Figure 3:**
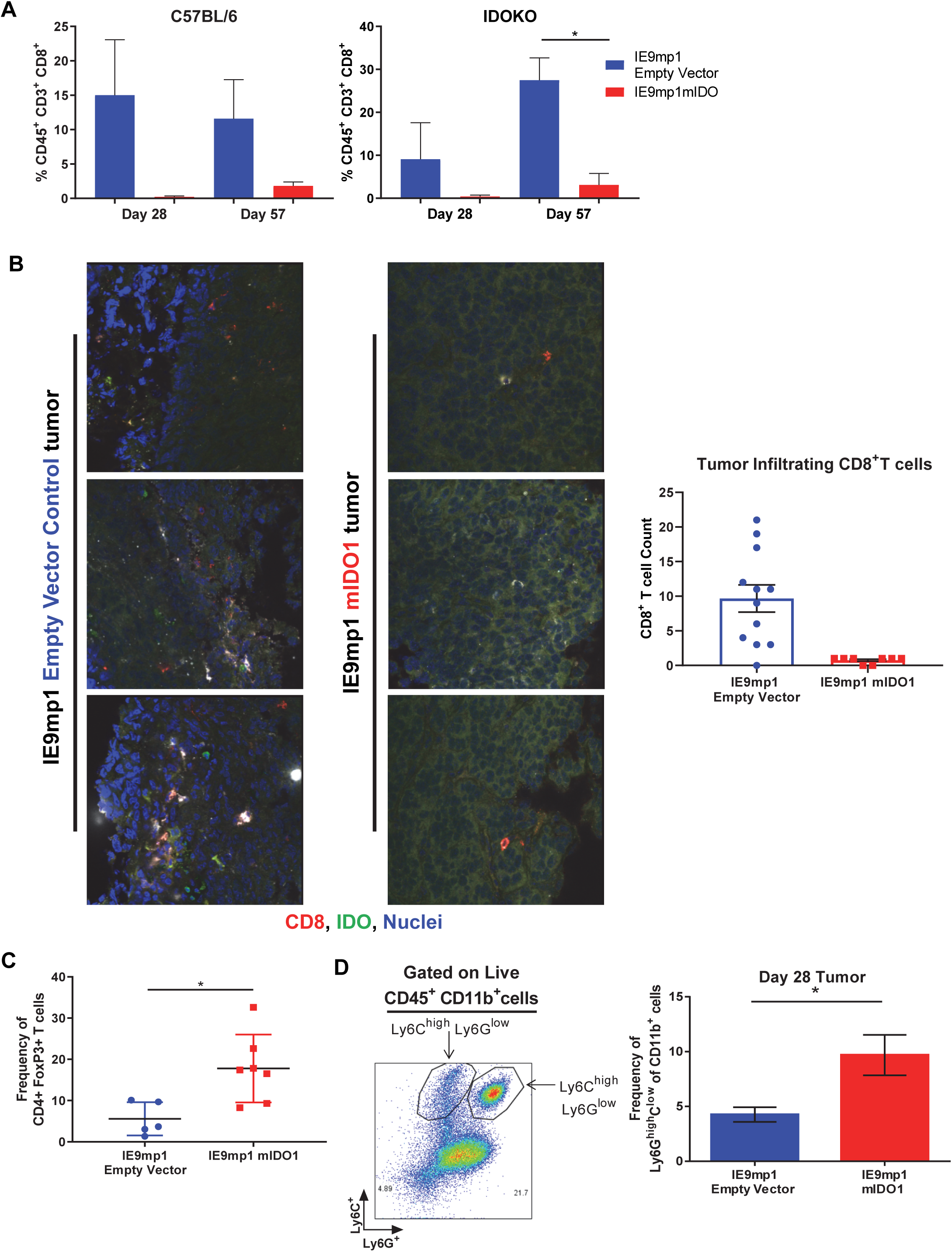
Expression of IDO1 leads to poor tumor infiltration by CD8+ T cells and increased infiltration of suppressive immune cells in the ovarian tumor microenvironment. **(A)** Frequency of CD8^+^ TILs on Days 28 and 57 from WT C57BL/6 (left panel n=6) and IDOKO (right panel n=6) mice challenged i.p. with 1×10^7^ IE9mp1-mIDO1 or IE9mp1-EV tumor cells. **(B)** Immunostained IE9mp1-mIDO1 and IE9mp1-EV tumor from WT C57BL/6 mice on Day 48 with anti-IDO1 (green), CD8^+^ T cell (red), and DAPI (blue). Quantification of CD8^+^ T cells from 3 field images per tissue slide, (n=11 IE9mp1-EV and n=8 IE9mp1-mIDO1). **(C)** Flow cytometry analysis of tumor ascites for CD4^+^CD25^+^FoxP3^+^ cell frequency in IE9mp1-mIDO1 (n=7) and IE9mp1-EV (n=5) tumor-bearing WT C57BL/6 mice on Day 48. **(D)** Frequency of Ly6G^high^Ly6C^low^ CD11b^+^ cell IE9mp1-EV (n=5) or IE9mp1-mIDO1 (n=5) tumor-bearing WT C57BL/6 mice on Day 28. ∗p < 0.05, ∗∗p < 0.01, using Student’s t test **(B, C** and **D)**. The data represent means ± SEM of three independent experiments.

As IDO1 is known to induce Tregs (9, 11, 30), ovarian TME CD4^+^CD25^+^FoxP3^+^ Treg frequency was examined in IDO1-sufficient C57BL/6 mice. Treg frequency was significantly elevated by Day 48 in IE9mp1-mIDO1 tumor-bearing mice (Figure 3C) along with increased levels of CD11b^+^Ly6G^high^Ly6C^low^ myeloid cells (Figure 3D). To address potential mechanisms regulating the influx and retention of Tregs and myeloid cells in the ovarian TME, chemokines were measured in cell-free tumor ascites fluid from C57BL/6 mice (31). The Treg-attractant MIP-1β/CCL4 (32) was significantly increased in IDO1-expressing tumors (Supplemental Figure 3A). Similarly, IDO1-expressing tumors exhibited significant increases in monocyte/macrophage attracting MCP-3/CCL7, eotaxin/CCL11, MCP-1/CCL2, and G-CSF (Supplemental Figures 3B-E) (33, 34). These results indicate that tumoral IDO1 expression regulates the ovarian TME chemokine signature for enhanced recruitment of immunosuppressive cells resulting in reduced CD8^+^ TIL frequency.

### Tumoral IDO1 mediates CD8^+^ TIL gene expression changes

The impact of IDO1 expression on CD8^+^ TIL frequency via alteration of the ovarian TME metabolic and cellular profile prompted us to investigate the consequences of tumoral IDO1 expression on gene expression changes of CD8^+^ TILs. Transcriptome analysis of IDO1-sufficient and IDO1-knockout CD8^+^ TILs isolated from IE9mp1-mIDO1 and IE9mp1-EV expressing tumors identified 5,163 differentially expressed genes (FDR<0.1) unique to IDOKO TILs and 4 differentially expressed genes unique to WT mice. We utilized the blood transcriptome modules (BTM) approach (35) to further investigate the transcriptomic changes. This approach is based on large-scale data integration of gene networks from over 30,000 transcriptomes in more than 500 studies, to generate over 300 context specific modules in the reconstructed network, providing a deeper insight than looking at individual genes. Unsupervised PCA revealed different gene expression patterns in CD8^+^ TILs mediated by expression or absence of IDO1 expression in host cells or tumor cells (Figure 4A). By examining the overall expression of BTMs, we observed separation of gene signatures into two groups by hierarchical clustering (Figure 4B). Cluster 1 (upper) comprises 103 gene modules downregulated in IDOKO CD8^+^ TILs present in IDO1-expressing tumors. Notably, the downregulated genes within Cluster 1 are related to the regulation of antigen presentation and immune responses, T cell activation and signaling, and CD28 co-stimulation (Supplemental Table 2). Pathway Enrichment Analysis further demonstrated that downregulated genes in Cluster 1 play a role in TCR signaling, the adaptive immune system, and oxidative phosphorylation in lymphocytes (Figure 4C). Conversely, the 244 gene modules with upregulated gene expression in IDOKO CD8^+^ TILs from IE9mp1-mIDO1 tumors in Cluster 2 (lower) (Figure 4B) were found to be enriched in G protein-coupled receptors (GPCR) downstream signaling (Supplemental Figure 4). GPCRs regulate T cell chemotaxis (36) and have been shown to play a crucial role in tumor progression and metastasis (37).

**Figure 4:**
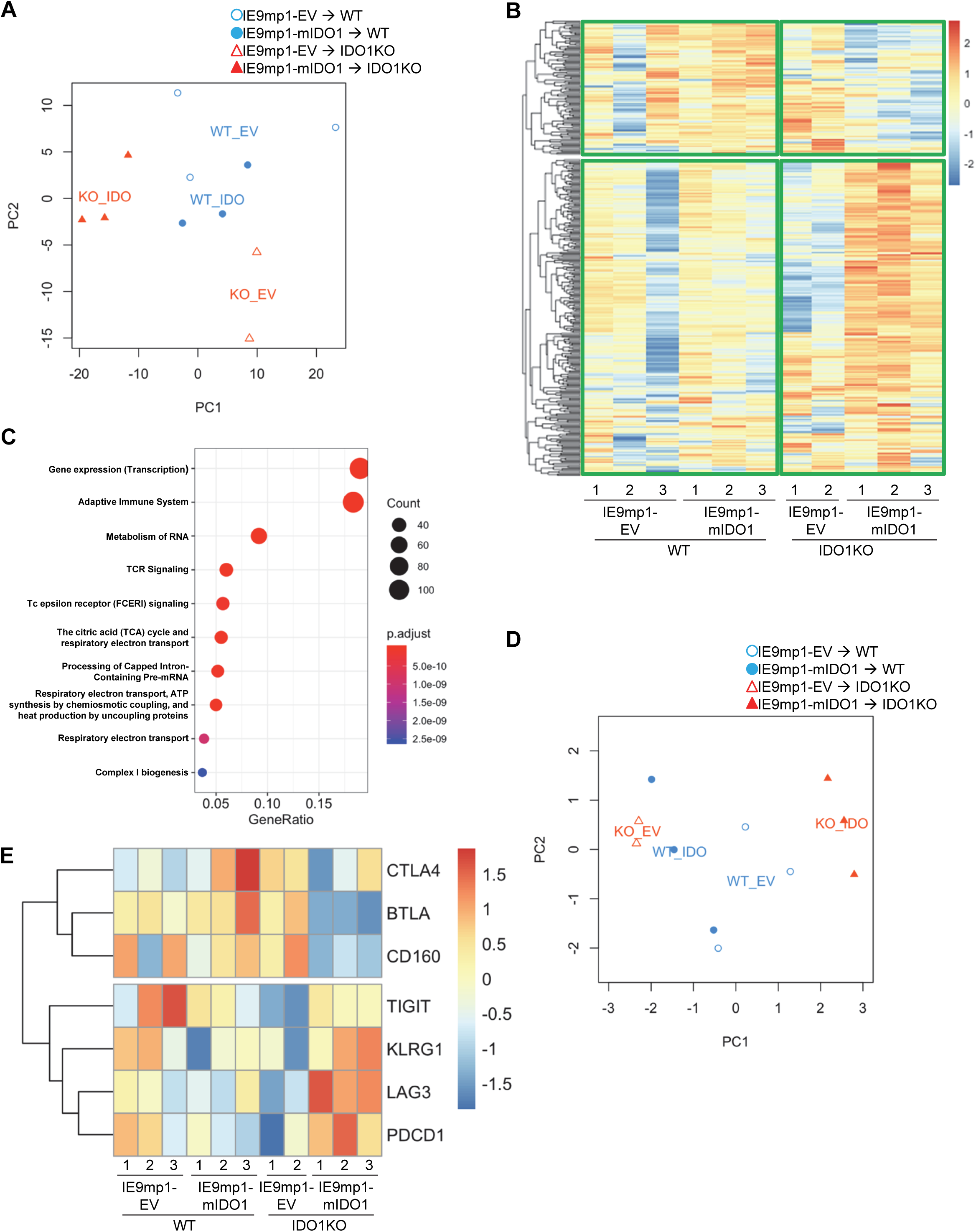
Tumoral IDO1 mediates gene expression changes in tumor infiltrating T cells. **(A)** PCA scatterplot on first two components shows transcriptional gene signature groups in CD8^+^ TILs from IE9mp1-mIDO1 (solid) and IE9mp1-EV (open) i.p. tumor-bearing WT C57BL/6 (circle) and IDOKO (triangle) mice at Day 48. **(B)** Heat map shows the blood transcriptome modules (BTMs) activity grouped into two clusters by hierarchical clustering. The measurements were obtained from four experimental groups, WT C57BL/6 mice – IE9mp1-mIDO1 tumor, WT C57BL/6 mice – IE9mp1-EV tumor, IDOKO mice – IE9mp1-mIDO1 tumor, and IDOKO mice – IE9mp1-EV tumor at Day 48. **(C)** Pathway Enrichment Analysis for Cluster 1 genes from the heat map. Gene Ratio shows difference in the number of genes enriched by the total number of genes in each pathway. Circles represent the number of genes. Color scale shows adjusted p-values. **(D)** PCA scatterplot of the change in inhibitory receptor gene expression in CD8^+^ TILs from IE9mp1-mIDO1 (solid) and IE9mp1-EV tumor (open) tumor-bearing WT C57BL/6 (blue circle) and IDOKO mice (red triangle) mice at Day 48. **(E)** Heat map of the change in inhibitory receptor gene expression in CD8^+^ TILs in tumors from the four experimental groups at Day 48.

The composition of gene modules in Cluster 1 suggests that tumoral IDO1 expression suppresses CD8^+^ TIL function, in particular focusing on regulators of T cell exhaustion and dysfunction (Supplemental Table 2). Consistent with the BTM analysis, PCA identified differences in expression of immune inhibitory receptors in CD8^+^ TILs between the IE9mp1-mIDO1 (solid) and IE9mp1-EV (open) tumor-bearing WT C57BL/6 (blue circle) and IDOKO (red triangle) mice (Figure 4D). Interestingly, the differences were more notable in IDOKO mice. Differential expression analysis on individual inhibitory genes showed that when IDO1 is expressed by the tumor, *PDCD1* (encoding PD-1), *LAG3*, and *KLRG1* were significantly upregulated on CD8^+^ TILs, while *CD160* and *BTLA* were significantly downregulated (p<0.05) in the IDOKO mice group (Figure 4E). Notably, *PDCD1* had the greatest fold-change (IE9mp1-mIDO1 vs. IE9mp1-EV) amongst the inhibitory genes analyzed in CD8^+^ TILs from IDOKO mice (Supplemental Table 2). Taken together, the data demonstrated that tumoral IDO1 led to the upregulation in expression of a set of inhibitory receptors on CD8^+^ TILs.

### Kynurenine mediates induction of inhibitory receptors on CD8^+^ T cells

To assess whether host- or tumor-derived IDO1 expression leads to upregulation of inhibitory receptors, we phenotyped CD8^+^ TILs isolated from WT and IDOKO mice bearing either IE9mp1-mIDO1 or IE9mp1-EV tumors. In both IDO1-sufficient and IDO1-knockout TILs, tumoral IDO1 expression significantly increased the frequency of PD-1^+^ CD8^+^ TIL by Day 48 compared with those from IE9mp1-EV challenged mice (Figure 5A). Notably, in the absence of host- and tumor-derived IDO1 this effect was abrogated and there was no significant upregulation of PD-1 on CD8^+^ TIL at the same time point (Day 48) in IE9mp1-EV tumor-bearing IDOKO mice, indicating that regardless of the cellular source of IDO1 expression, presence of IDO1 enzyme activity contributes to PD-1 upregulation on ovarian CD8^+^ TIL.

**Figure 5:**
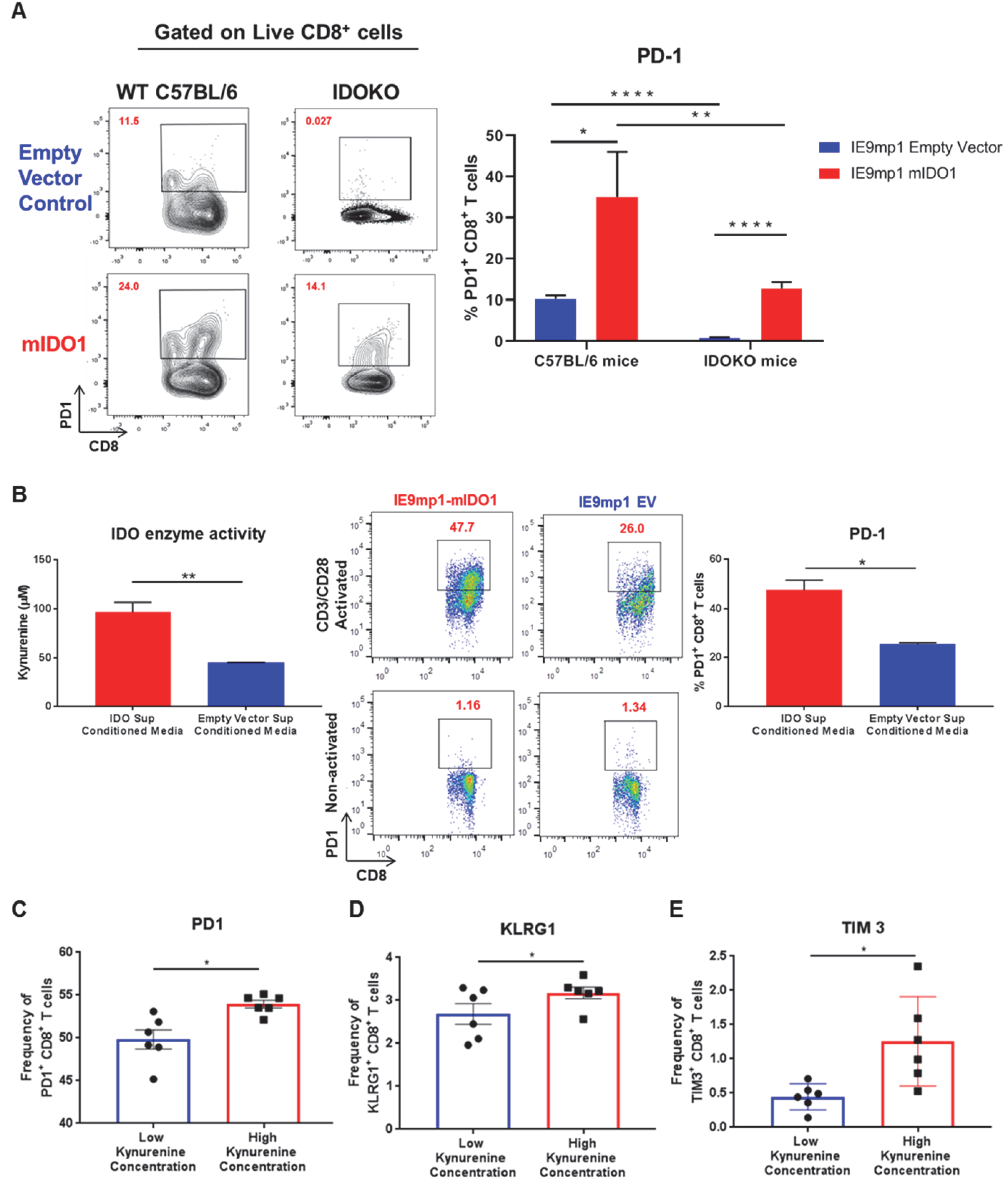
Kynurenine mediates induction of inhibitory receptor on CD8^+^ T cells. **(A)** Frequency of PD-1^+^ CD8^+^ T cells in tumor ascites from WT C57BL/6 or IDOKO mice challenged i.p. with 1×10^7^ IE9mp1-EV (n=5) or IE9mp1-mIDO1 (n=6) tumor cells on Day 48, as determined by flow cytometric analysis. **(B, left)** Kynurenine concentrations measured in IE9mp1-EV and IE9mp1-mIDO1 tumor cell culture supernatant by colorimetric assay. **(B, right)** Frequency of CD8^+^PD-1^+^ T cells, from WT C57BL/6 mice spleens, co-cultured with 1μg/mL anti-CD3/CD28 in IE9mp1-mIDO1 or IE9mp1-EV tumor cell culture supernatant for 48 hr. CD8^+^PD-1^+^ T cell frequency was determined by flow cytometric analysis. **(C)** PD-1, **(D)** KLRG1, and **(E)** TIM3 expression analyzed by flow cytometric analysis on lymphocytes from WT C57BL/6 mice activated with 1μg/mL anti-CD3/CD28 and with treated kynurenine. ∗p < 0.05, ∗∗p < 0.01, ∗∗∗∗p < 0.0001, by Student’s t test **(A, B, C, D** and **E)**. The data represent means ± SEM of three independent experiments performed in triplicate.

The contribution of IDO1-induced Kyn generation by tumor cells on PD-1 expression was evaluated using anti-CD3/anti-CD28 activated CD8^+^ lymphocytes. IE9mp1-mIDO1 cell culture supernatant allowed for a controlled system where IDO1-mediated Trp catabolism into Kyn could be quantified (Figure 5B, left). CD8^+^ T cells significantly upregulated PD-1 expression in the presence of elevated tumor-produced Kyn (Figure 5B, right). Activation of CD8^+^ T cells in varying Kyn concentrations resulted in upregulation of not only PD-1 (Figure 5C), but also additional inhibitory receptors such as KLRG1 (Figure 5D) and TIM3 (Figure 5E), although the effect was more pronounced at higher Kyn concentrations. Taken together, these results demonstrated that kynurenine contributed to the upregulation of co-inhibitory receptors expression on CD8^+^ T cells *in vivo* and *in vitro*.

### PD-1 gene contains putative AHR binding sites

Upon activation by its natural endogenous agonist, Kyn, AHR translocates into the nucleus and regulates target gene expression in T cells (10, 11), including a tumor-promoting role via suppression of anti-tumor immunity (24). To account for this observation, we hypothesized that the genes encoding T cell inhibitory receptors contain AHR binding sites responsible for transcriptional regulation upon sequential Kyn ligation and AHR activation. Therefore, we performed computational analysis of promoter regions of T cell inhibitory receptor genes for the consensus AHR xenobiotic response elements (XRE) binding motifs (Figure 6A). We identified multiple AHR binding sites in the upstream gene promoter region of murine (Figure 6B) and human *PDCD1* (PD-1) gene (Supplemental Figures 5 and 6). AHR binding sites were also present in the upstream promoter regions of additional inhibitory receptors such as LAG3, TIM3, KLRG1, CTLA4, BTLA, 2B4, CD160 and TIGIT.

**Figure 6:**
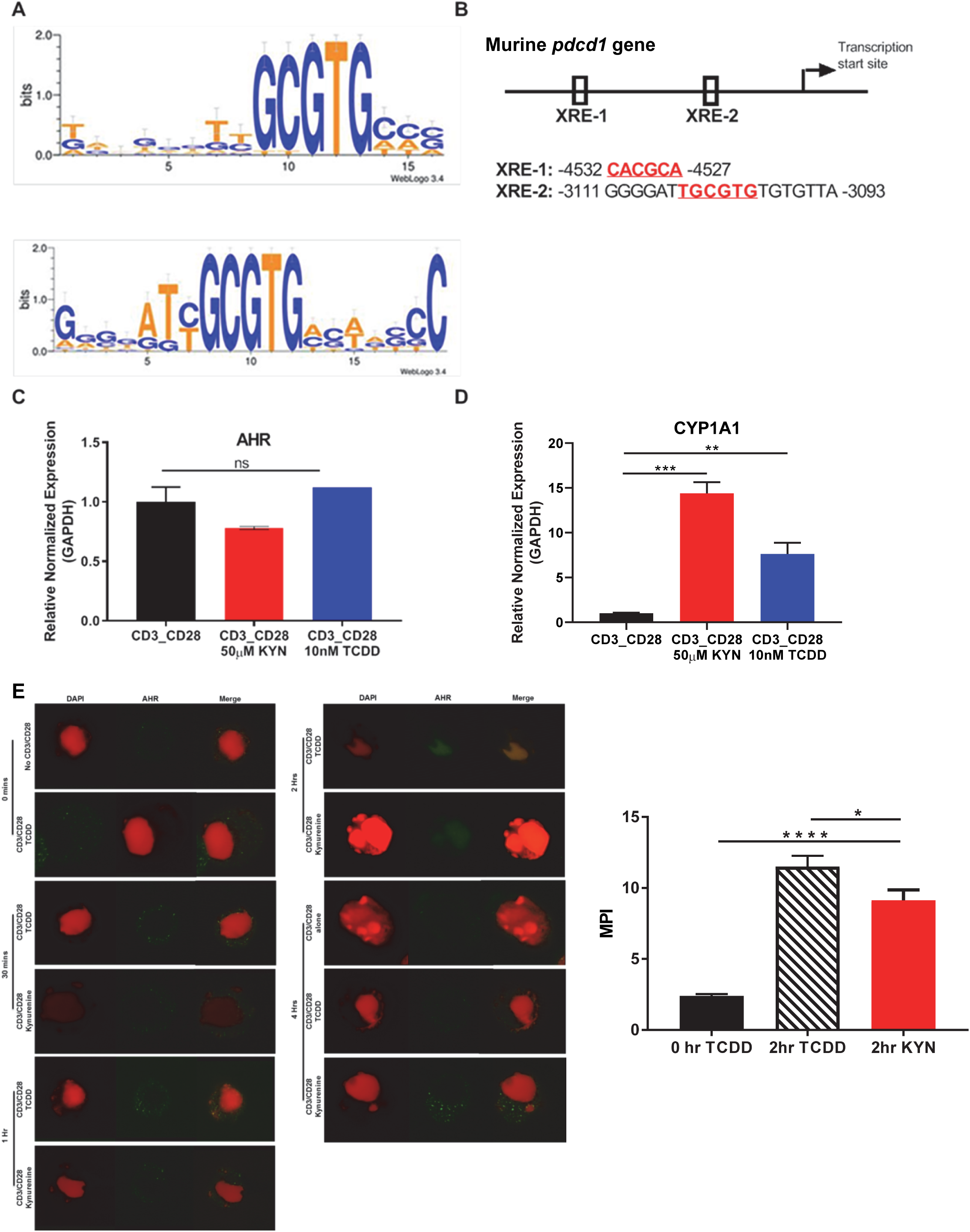
PD-1 gene contains putative aryl hydrocarbon receptor binding sites. **(A)** Motif Logo depicting the position specific weight matrices of the AHR binding site (xenobiotic response element (XRE)). **(B)** Schematic representation of the identified AHR binding site in the *PDCD1* promoter. **(C)** Anti-CD3/CD28 activated CD8^+^ T cells from spleens of WT C57BL/6 mice were treated with IL-2 (50U/mL), KYN (50µM) or TCDD (10nM) for 6 days. *AHR* and **(D)** *cyp1a1* mRNA was determined. **(E)** AHR nuclear translocation was measured in anti-CD3/CD28 bead-activated CD8^+^ T cells from spleens of WT C57BL/6 mice treated with KYN (50µM) or TCDD (10nM) for 0 mins, 30 mins, 1 hr, 2 hrs and 4 hrs. CD8^+^ T cells were immunostained for AHR (green) and DAPI (red). Mean fluorescence intensity (MFI) of AHR nuclear translocation was calculated (bar graph). ∗p < 0.05, ∗∗p < 0.01, ∗∗∗p < 0.001, ∗∗∗∗p < 0.0001, ns: not significant, by Student’s t test **(D** and **E)**. The data represent means ± SEM of three independent experiments performed in triplicate.

To address the mechanism by which Kyn mediates PD-1 expression, we next evaluated AHR expression in activated CD8^+^ T cells. AHR gene expression in T cells was not altered by their activation in the presence of Kyn or another well characterized exogenous AHR ligand, 2,3,7,8-tetrachlorodibenzo-p-dioxin (TCDD) (Figure 6C) (38). Although Kyn ligation did not impact AHR expression in T cells, it did result in AHR target gene transcription, specifically the cytochrome P40 enzyme, CYP1A1 (Figure 6D) (39), confirming Kyn-mediated AHR activation. As CYP1A1 expression requires nuclear translocation of the active AHR-Kyn complex (40, 41), we evaluated the kinetics of Kyn-mediated AHR nuclear translocation in activated CD8^+^ T cells. At 2 hrs post-activation in the presence of Kyn and TCDD, AHR was observed in the nucleus (Figure 6E, green) and thus provides a mechanism by which AHR directly mediates PD-1 expression in CD8^+^ T cells.

### Kynurenine permits genome-wide chromatin accessibility in regulatory regions of PD-1 gene

As Kyn-mediated upregulation of inhibitory receptors requires AHR interaction with XRE sequences in the promoter regions, we next evaluated how Kyn treatment alters the dynamics of CD8^+^ T cell genome-wide chromatin accessibility by ATAC-seq (42). Increases of chromatin accessibilities in Kyn-treated activated CD8^+^ T cells were observed for two regulatory elements of the PDCD1 (Figure 7A) and LAG3 (Supplemental Figure 7) genes, indicating that Kyn treatment mediates the extent of chromatin accessibility for key T cell inhibitory receptors leading to their upregulation. Moreover, these data also suggest an epigenetic role for IDO1 in regulating the transcriptional activity of CD8^+^ TILs.

**Figure 7:**
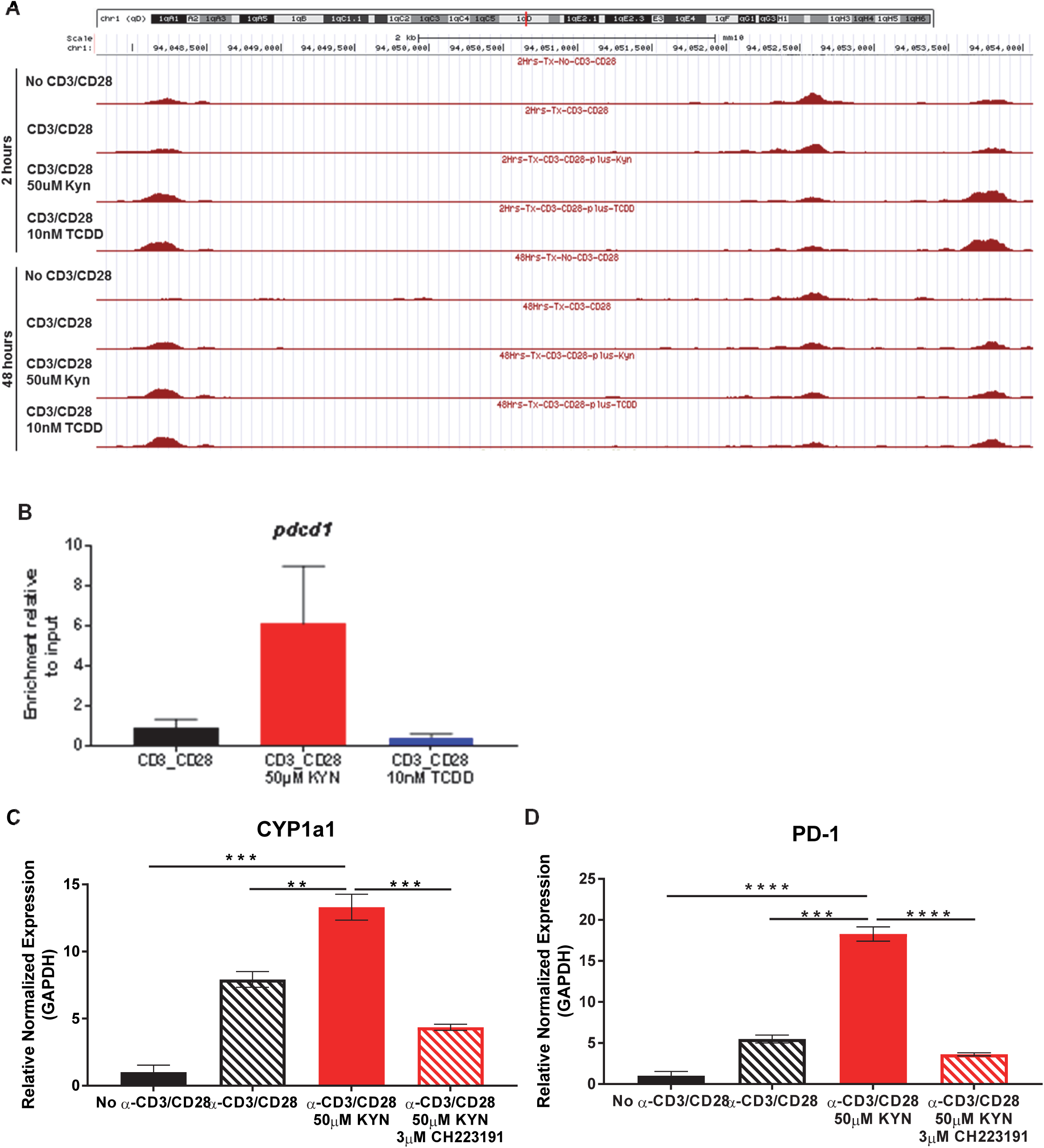
Kynurenine induces PD-1 expression on CD8^+^ T cells in an AHR-dependent manner. **(A)** UCSC Genome Browser plot of open chromatin accessible regions in two DNA regulatory elements in the PD-1 gene identified by ATAC-seq analysis of CD8^+^ T cells, from WT C57BL/6 mice, activated by anti-CD3/CD28 and treated with KYN (50µM) or TCDD (10nM) for 2 and 48 hrs. **(B)** ChIP-qPCR PDCD1 analysis in anti-CD3/CD28 bead-activated and KYN (50µM) treated CD8^+^ T cells for 2 hrs from spleens of WT C57BL/6 mice with AHR antibody and *pdcd1* promoter primers. **(C)** Anti-CD3/CD28 activated CD8^+^ T cells from spleens of WT C57BL/6 mice were treated with KYN (50µM) and CH223191 (3μM) for 6 days. *Cyp1a1* and **(D)** *PD-1* mRNA was determined. ∗∗p < 0.01, ∗∗∗p < 0.001, ∗∗∗∗p < 0.0001 by Student’s t test **(B, C** and **D)**. The data represent means ± SEM of three independent experiments performed in triplicate **(B, C** and **D)**.

### Kyn induces PD-1 expression on CD8^+^ T cells in an AHR-dependent manner

Direct evidence for Kyn-mediated AHR binding to XRE motifs in the PD-1 promoter was evaluated by chromatin immunoprecipitation of activated CD8^+^ T cells (Supplemental Figure 8). ChIP-qPCR using anti-AHR bound to genomic DNA confirmed enrichment of AHR binding to XRE motifs within the murine PDCD1 gene promoter region by Kyn treatment but not by TCDD (Figure 7B). Furthermore, ChIP-qPCR verified PD-1 gene expression was upregulated in the presence of Kyn. The requirement for AHR-mediated PD-1 expression following Kyn treatment was tested by utilizing the AHR antagonist CH223191 (43). In addition to inhibiting its downstream target gene CYP1A1, AHR antagonism severely diminished Kyn-mediated PD-1 upregulation on CD8^+^ T cells (Figure 7C and D). These data confirm that AHR interaction with responsive elements in the PD-1 promoter facilitates its transcription upon treatment with Kyn and suggests an approach in which AHR antagonism can be coupled with IDO1 blockade to synergistically prevent PD-1 mediated T cell dysfunction.

## Discussion

Tryptophan catabolism by IDO1 has been firmly established as a powerful mechanism of innate and adaptive immune tolerance in EOC and other solid tumors (14, 15, 17, 44). However, efforts to target this pathway in the clinic has met with limited success (18, 19), probably because the full biologic consequences of IDO1 on the TME remain incompletely characterized. In this study, we fill a major gap in knowledge by demonstrating that (i) the metabolic re-wiring of the TME by IDO1 is profound and goes beyond Kyn accumulation by also affecting nicotinate/nicotinamide and purine metabolism; (ii) IDO1 leads to significant quantitative (frequency) and qualitative (gene expression) effects on CD8^+^ TILs, including upregulation of inhibitory receptors; (iii) Kyn alters chromatin accessibility in regulatory regions of inhibitory receptors leading to their upregulated expression; and (iv) AHR binding to consensus XRE motifs is the mechanism by which IDO1 activity mediates T cell dysfunction. These results support the design of strategies to target the IDO1 and AHR pathways for improving anti-tumor immunity in EOC.

To study the consequences of the metabolic microenvironment imposed by IDO1, we first investigated quantitative and qualitative changes in CD8^+^ T cell infiltration. In addition to the effect of IDO1 in reducing intratumoral accumulation of CD8^+^ TILs, the cytokine milieu of the TME was noted to favor the recruitment of immunosuppressive Tregs and myeloid cells. Moreover, there were distinct changes in expression of clusters of genes related to T cell activation, co-stimulation and signaling. This led to a focus on T cell inhibitory receptors, which indicated upregulation of several inhibitory receptors including PD-1, LAG3, and KLRG1. Tumoral IDO1-induced Kyn was sufficient to upregulate PD-1 on CD8^+^ T cells *in vivo* and *in vitro*. Moreover, we established AHR activation by Kyn as the mechanism by which CD8^+^ T cells acquired an exhausted phenotype of PD-1 expression.

Although our bioinformatics analyses identified AHR XRE binding sites in the promoter region of several human and murine checkpoint receptor genes, chromatin accessibility by ATAC-seq indicated the potential for AHR transcription factor occupancy in PD-1 and LAG3 of inhibitory receptor genes. Since ATAC-seq alone cannot decide which transcription factor binds to accessible chromatin (42), we utilized ChIP-qPCR and demonstrated specific binding of AHR to DNA in the PD-1 gene. Thus, Kyn increased the accessibility of AHR to XRE sequences in the PD-1 gene promoter of CD8^+^ T cells. Further support for an AHR-dependent mechanism for kynurenine upregulation of PD-1 gene expression is provided by abrogation of gene expression in the presence of the AHR antagonist CH223191. These findings establish that kynurenine activation of AHR is a critical mechanism by which IDO1 impacts the upregulation of checkpoint receptors on CD8^+^ T cells in the ovarian TME. Although PD-1 expression was shown to be mediated by non-physiological concentrations of Kyn in a recent study (45), the downstream transcriptional events that regulate this dysfunctional TIL phenotype were not identified. Our study provides a targetable metabolic mechanism by which IDO1 regulates checkpoint receptor expression on CD8^+^ TILs through the transcriptional regulation of AHR activation upon Kyn generation from host and tumor cellular sources.

A notable limitation of the present study is that while we have uncovered alterations in metabolic pathways beyond kynurenine, their contribution to induction of T cell inhibitory receptors or other mechanisms of immune suppression were not fully examined. For example, NAD is able to regulate CD4 T cell differentiation and promotes IL-10 and TGF-β1 production by Th1 and Th17 cells, respectively (46). In addition, it will be important to confirm whether AHR mediated upregulation of inhibitory T cell receptors is a general mechanism for other metabolites along the kynurenine pathway. Moreover, while murine studies can yield valuable insights into AHR function, there are differences between the human and mouse AHR, such as differences in the affinity to ligand activation (47).

Despite these limitations, the current study sheds some light on potential reasons for the limited efficacy of the combination of IDO1 and PD-1 blockade in clinical trials (20–22). First, Kyn activation of AHR may be a major resistance mechanism via upregulation of several immune checkpoint receptors, with the implication that PD-1 blockade monotherapy may be insufficient to derive benefit from IDO-1 blockade. In particular, we demonstrate that LAG3 gene expression on CD8^+^ TILs is similarly impacted by Kyn, and therefore LAG3 may serve as a compensatory escape pathway when the PD-1/PD-L1 pathway is blocked. Second, AHR antagonists have been identified and examined as potential therapeutic tools to study the role of AHR in tumorigenesis (38). Preclinical studies in multiple myeloma, suggests that therapeutic targeting of the AHR, with an inhibitor such as the FDA-approved clofazimine (48), may improve clinical outcomes.

Altogether, this study demonstrates that IDO1-induced Kyn activates AHR nuclear translocation where its direct binding to XRE motifs in the PD-1 gene promoter (Supplemental Figure 9) results in T cell dysfunction. Strategies to concomitantly target the IDO1 and AHR pathways may overcome immune suppression and enhance anti-tumor immunity in EOC and other solid tumors.

## Methods

### The Cancer Genome Atlas (TCGA) data analysis

Gene expression data were downloaded from cBioportal. Immune and IDO signature scores were calculated using the sum of the gene expression selected genes (Supplemental Table 1) divided by the square root of the number of genes. Patients were then divided in four groups: TIL^High^/IDO^Low^, TIL^Low^/IDO^Low^, TIL^Low^/IDO^High^, and TIL^High^/IDO^High^ based on immune and IDO score values. Survival analysis across groups was calculated using clinical information available on cBioportal using log rank test at a significance threshold of 0.05.

### Animals

Female and male WT C57BL/6 mice were purchased from The Jackson Laboratory (Bar Harbor, ME) and bred in our facility (Roswell Park Comprehensive Cancer Center) according to an approved protocol. IDO1-knockout (IDOKO) mice were purchased from Jackson Laboratory (Bar Harbor, ME). All animals were maintained in the Laboratory Animal Shared Resource under specific pathogen-free conditions. All animal experiments were carried out according to protocol guidelines approved by the Institute Animal Care and Use Committee (IACUC) of Roswell Park Comprehensive Cancer Center (Buffalo, NY).

### Cell Lines

The murine IDO1 (mIDO1) gene was overexpressed in mouse ovarian surface epithelial cell (MOSEC) lines by retroviral transduction. Full-length mIDO1 gene was PCR amplified from cDNA of IFN-γ-treated ID8 cell line (26) and inserted into the first cloning site in a retrovirus backbone vector (pQCXIX, Clontech-TaKaRa). A fusion gene of codon-optimized Luciferase (Luc2) and tandem-dimeric Tomato (tdT) gene was PCR amplified from the pcDNA3.1(+)Luc2=tdT (a gift from Christopher Contag (Addgene plasmid # 32904))(49) and inserted into the second cloning site. A control empty vector was constructed by inserting Luc2=tdT gene alone. The retroviral transfer vector was co-transfected with pVSV-G retroviral envelope-expressing plasmid into the GP2-293 cell line (Clontech-TaKaRa) using Lipofectamine 2000 reagent (Invitrogen) to produce retrovirus supernatant. Parental ID8 (26), IE9 (50) or IE9mp1 (25) tumor cell lines were transduced by retroviral vectors in the presence of 8mg/ml polybrene (Sigma-Aldrich). Stable cell lines were established after flow cytometry cell sorting of tdT-expressing cells using a FACSAria II (BD Biosciences). Expression of tdT in established cell lines was periodically monitored by flow-cytometry and confirmed that >99% cells expressed tdT before experiments. All cell lines were cultured in complete culture medium: RPMI1640 (Corning Cellgro®) supplemented with 10% fetal bovine serum (VWR), 1% sodium pyruvate (100mM), 1% L-glutamine (200mM), 1% MEM nonessential amino acid (100×), 1% penicillin/streptomycin (100×), 2.5% Hepes, and 0.1% beta-2-mercaptoethanol (50mM) in an incubator at 37°C and 5% CO_2_.

### Tumor challenge and measurement

WT C57BL/6 and IDOKO mice were challenged intraperitoneally (i.p.) with 1×10^7^ IE9mp1-mIDO1 or IE9mp1-EV tumor cells in a final volume of 500μl Dulbecco’s PBS (Corning Cellgro®). Tumor progression and the amount of tumor burden were monitored by measuring abdominal distension to track the accumulation of peritoneal ascites formation. Mice were euthanized by CO_2_ asphyxiation and/or cervical dislocation when the abdominal circumference of i.p. tumors reached a 50% girth increase and/or upon detection of declining health conditions as described in our standard operating procedure for Body Scoring, according to IACUC guidelines.

### IDO enzyme activity measurement

Kynurenine was measured in cell culture supernatants by incubating the sample with a 30% w/v Trichloroacetic acid (Sigma-Aldrich) solution prepared in water, and Ehrlich reagent (2% PDAB) prepared fresh for each assay by dissolving p-dimethylaminobenzaldehyde (PDAB) (Sigma-Aldrich) in acetic acid. The colorimetric reaction was measured using a microplate reader at OD 490nm. The OD values were measured and calculated against standard dilution curve of L-kynurenine (Sigma-Aldrich).

### Flow Cytometry antibodies and reagents

LIVE/DEAD™ Fixable Yellow Dead Cell Stain Kit, for 405nm excitation was purchased from Thermo Fischer Scientific. Mouse-specific FITC-anti-CD45 (30-F11), eFluor450-anti-CD11b (M1/70), APC-anti-TIM3 (8B.2C12), APC-anti-CD25 (PC61.5), PerCP-eFluor710-anti-CD8α (53-6.7), PE-anti-Foxp3 (FJK-16s), and PE-Cy7-anti-CD4 (GK1.5) were purchased from eBiosciences. BV421-anti-PD-1 (29F.1A12) was purchased from BioLegend. Anti-CD16/CD32 FcBlock (2.4G2), PE-Cy7-anti-Ly6G/Ly6C (RB6-8C5), BV450-anti-CD45 (30-F11), FITC-anti-CD8α (53-6.7), PerCP-Cy5.5-anti-Ly6G/Ly6C (RB6-8C5), PerCP-anti-CD8α (53-6.7), PE-anti-CD3ε (145-2C11), PE-anti-CD11b (M1/70), and PerCP-anti-CD4 (RM4-5) were purchased from BD Biosciences. Foxp3/Transcription Factor Fixation and Permeabilization Concentrate and Diluent kit (eBiosciences) was used for intracellular Foxp3 staining. Flow cytometry data were acquired using a BD Biosciences LSRII flow cytometer and BD FACSDiva software, and analyzed using FlowJo v10 software (TreeStar).

### RNA isolation and quantitative real-time polymerase chain reaction (PCR) analysis

Total RNA was isolated using the RNeasy mini kit (Qiagen). cDNA was prepared using iScript^TM^ cDNA Synthesis kit (BioRad) and reverse transcription was carried out on the T100^TM^ Thermal Cycler (BioRad). Quantitative real-time PCR was performed using iQ^TM^ SYBR Green Supermix (BioRad) on the C1000 Touch Thermal Cycler, CFX96^TM^ Real-Time System (BioRad). Primer sequences used include *IDO1* forward 5’-TCTGCCTGTGCTGATTGA-3’, reverse, 5’-CTGTAACCTGTGTCCTCTCA-3’; *AHR* forward 5’-CCACTGACGGATGAAGGA-3’, reverse, 5’-ATCTCGTACAACACAGCCTCT-3’; *CYP1A1* forward 5’-GACACAGTGATTGGCAGAG-3’, reverse, 5’-GAAGGTCTCCAGAATGAAGG-3’; *PDCD1* forward 5’-CTCGGCCATGGGACGTAGGG-3’, reverse, 5’-GGGTCTGCAGCATGCTAATGGCTG-3’; and *GAPDH* forward 5’-GCCTTCCGTGTTCCTACCC-3’, reverse, 5’-CAGTGGGCCCTCAGATGC-3’. PCR data were analyzed using CFX Manager 3.1 (BioRad). Experiments were performed in triplicates.

### Purification of CD8^+^ T cells from splenocytes

CD8^+^ T cells were isolated from spleens of WT C57BL/6 mice by negative selection using the EasySep™ Mouse CD8^+^ T cell isolation kit (Stem Cell Technologies) according to the manufacturer’s protocol and cultured in complete media. Isolated CD8^+^ T cells were activated with either plate-bound (1μg/mL) anti-CD3 and soluble (1μg/mL) anti-CD28, or Dynabeads™ Mouse T-Activator CD3/CD28 beads (Thermo Fischer Scientific) and co-cultured in the presence or absence of 50U/mL IL-2, 50μM kynurenine (Sigma Aldrich), 3μM CH223191 (Sigma-Aldrich) or 10nM 2,3,7,8-tetrachlorodibenzo-p-dioxin (TCDD, Thomas Scientific).

### Isolation of CD8^+^ T cells from tumor

Liberase Thermolysin Medium Research Grade solution (Roche) was used for the digestion of IE9mp1-mIDO and IE9mp1-EV tumor tissues harvested from WT C57BL/6 and IDOKO mice. Following digestion, direct *ex vivo* analysis by flow cytometry was performed in order to phenotype immune cell infiltration into the tumor. To lyse red blood cells in hemorrhagic tumor ascites fluids in order to perform phenotypic immune cell analysis, ACK lysis buffer was used. Ammonium-Chloride-Potassium (ACK) lysing buffer was added to ascites fluid and incubated at room temperature, then washed with 1× PBS before staining for surface and intracellular markers.

### Cytokine and chemokine analysis

Cell-free tumor ascites fluid from IE9mp1-EV or IE9mp1-mIDO1 tumor-bearing WT C57BL/6 mice at Days 28 and 48 were harvest and analyzed for cytokines and chemokine levels using the 36-Plex Mouse ProcartaPlex^TM^ Panel 1A Multiplex Luminex immunoassay kit (eBioscience). Data were collected using the Luminex-100 system and analyzed using StarStation Version 1.8 software (Applied Cytometry).

### Confocal microscopy

IE9mp1-mIDO1 or IE9mp1-Empty Vector tumor excised from WT C57BL/6 mice on Day 48 of tumor progression, or naïve CD8^+^ T cells from WT C57BL/6 mice activated with anti-CD3/anti-CD28 and treated with kynurenine (50μM) or TCDD (10nM) alone for 0 minutes, 30 minutes, 1 hour, 2 hours and 4 hours, were embedded in Tissue Freezing Medium (Ted Pella, Inc.) containing cryomolds and immediately frozen in 2-methyl-butane (Sigma-Aldrich). 5μm frozen sections of the tissues were made using the cryostat and layered on Superfrost™ Plus Slides (Thermo Scientific). The slides were fixed in 4% para-formaldehyde for 15 minutes, washed and blocked for 60 minutes at room temperature. The slides were then stained for 3 hours at room temperature with mouse-specific antibodies anti-IDO (D7Z7U, Cell Signaling Technologies), anti-CD8 (53-6.7, BioLegend), anti-PD-1 (J43, BD Biosciences), anti-AHR (W16012A, Biolegend), and 4’,6-Diamidino-2-Phenylindole (DAPI, Invitrogen). Confocal analyses of stained slides were performed using a TCS SP8 Laser Scanning Spectral Confocal Microscope (LEICA Microsystems). For enumeration of cells positive for the individual markers, photographs of 10 fields (at 63× magnification) of stained tumors or cells were taken and each field was counted using ImageJ 1.74v software (NIH). Mean cell counts of total 10 fields were plotted.

### Reagents and internal standards for High-performance liquid chromatography (HPLC)

Sources for reagents were: HPLC-grade acetonitrile and water (Burdick & Jackson); mass spectrometry-grade formic acid and ammonium acetate (Sigma-Aldrich); calibration solution containing multiple calibrants in a solution of acetonitrile, trifluroacetic acid, and water (Agilent Technologies); metabolite standards and internal standards, including N-acetyl Aspartic acid-d3, Tryptophan-15N2, Sarcosine-d3, Glutamic acid-d5, Thymine-d4, Gibberellic acid, Trans-Zeatine, Jasmonic acid, 15N Anthranilic acid, and Testosterone-d3 (Sigma-Aldrich).

### Sample preparation for mass spectrometry and metabolomics analysis

Metabolites were extracted from cell lines and the extraction procedure was previously described (51). Briefly, mouse liver tissue pool was used as quality controls in the extraction procedure. The extraction step started with the addition of 750µL ice-cold methanol:water (4:1) containing 20μL spiked internal standards to each tissue and quality control samples. Ice-cold chloroform and water were added in a 3:1 ratio for a final proportion of 2:4:3 water:methanol:chloroform. The organic (methanol and chloroform) and aqueous layers were collected, dried and resuspended with 500μL of 50:50 methanol:water. The extract was deproteinized using a 3kDa molecular filter (Millipore Corporation) and the filtrate was dried under vacuum (Gardiner). Prior to mass spectrometry, the dried extracts were re-suspended in 100μL of injection solvent composed of 1:1 water:methanol and were subjected to liquid chromatography-mass spectrometry. The injection volume was 10μL.

### LC-MS methods

For IDO pathway metabolites, ESI positive mode was used and analyzed using a 6495B triple quadrupole mass spectrometer (Agilent Technologies) coupled to a HPLC system (Agilent Technologies) via single reaction monitoring (SRM). Source parameters were: gas temperature-290°C; gas flow 14L/min; nebulizer 20psi; sheath gas temperature 350°C; sheath gas flow 12L/min; capillary 3000V positive and 3000V negative; nozzle voltage 1500V positive and 1500V negative. Approximately 8– 11 data points were acquired per detected metabolite. The HPLC column used was Zorbax eclipse XDB C-18, 1.8μm, 4.6×100mm (Agilent Technologies). Mobile phase A and B were 0.1% formic acid in water and acetonitrile respectively. Gradient used was: 0 min at 2% B; 2 min at 10% B, 12 min at 80% B, 18 min at 2% B followed by re-equilibration till end of the gradient 25 min to the initial starting condition (2% B). Flow rate used was 0.3mL/min. For purines and pyrimidines metabolites ESI positive mode was used and analyzed using a 6495B triple quadrupole mass spectrometer (Agilent Technologies) coupled to a HPLC system (Agilent Technologies) via single reaction monitoring (SRM). Source parameters, flow rate, and number of data points collected for purine/pyrimidines were similar to IDO pathway metabolites. A Waters X-bridge amide 3.5μm, 4.6×100mm column was used. Gradient used was: 0 min at 2% B; 6.50 min at 30% B; 7 min at 90% B; 12 min at 95%; 13 min at 2% B followed by re-equilibration till end of the gradient 20 min to the initial starting condition (2% B).

### Clonetech SMART-Seq v4 Ultra Low Input mRNASeq

Amplified cDNA was generated using the SMART-Seq v4 Ultra Low Input RNA kit (Clonetech). 1-1000 cells are collected in <9.5μL of PBS. The cells are lysed and first-strand cDNA was synthesized utilizing proprietary template switching oligos. Amplified ds cDNA was created by LD PCR using blocked PCR primers. The resulting ds cDNA was purified using AmpureXP beads (Beckman Coulter). Purified ds cDNA was quantitated by Qubit hs DNA (Thermo Fisher Scientific) and 1ng of material is used for subsequent library generation utilizing Nextera XT reagents (Illumina). Tagmentation was performed to shear the full-length cDNA to the appropriate size and incorporate partial adapter sequences. 12 cycles of PCR using Nextera XT primers complete the library and incorporate unique sample barcodes. The final libraries were purified using AmpureXP beads, and validated for appropriate size on a 4200 TapeStation D1000 Screentape (Agilent Technologies). The DNA libraries were quantitated using KAPA Biosystems qPCR kit, and were pooled together in an equimolar fashion, following experimental design criteria. The pool was denatured and diluted to 2.4pM with 1% PhiX control library added. The resulting pool was then loaded into the appropriate NextSeq Reagent cartridge for 75 cycle single-read sequencing and sequenced on a NextSeq500 following the manufacturer’s recommended protocol (Illumina).

### Assay for Transposase Accessible Chromatin sequencing (ATAC-seq)

50,000 isolated CD8^+^ T cells were activated with Dynabeads™ Mouse T-Activator CD3/CD28 beads (Thermo Fischer Scientific), and co-cultured in the presence of 50μM kynurenine (Sigma-Aldrich) or 10nM TCDD (Thomas Scientific) for 2 or 28 hours, and ATAC-seq DNA libraries were prepared as previously described (42, 52). The final libraries were purified using AmpureXP beads, and validated for appropriate size on a 4200 TapeStation D1000 Screentape (Agilent Technologies). The DNA libraries were quantitated using KAPA Biosystems qPCR kit, and were pooled together in an equimolar fashion, following experimental design criteria. Each pool was denatured and diluted to 350pM with 1% PhiX control library added. The resulting pool was then loaded into the appropriate NovaSeq Reagent cartridge for 100 cycle paired-end sequencing and sequenced on a NovaSeq6000 following the manufacturer’s recommended protocol (Illumina). ATAC-seq data in raw FASTQ format were processed uniformly through ENCODE ATAC-seq pipeline (version 1.4.0, https://github.com/ENCODE-DCC/atac-seq-pipeline). The sequences in FASTQ file were aligned to mouse genome version mm10 using BOWTIE2 in Paired-end mode. The mitochondria reads, the duplicated reads and the reads with mapping quality less than 30 were removed from downstream analysis. Then MACS2 (--extsize73 –shift 37) was used to call accessible regions and to generate genome-wide insertion sites profiles for visualization. The insertion sites profile shows number of insertion events found in certain genomic location normalized by sequencing depth by million reads. ATAC-seq data quality was controlled by estimating signal-to-noise ratio, transcription start sites (TSS) enrichment, and enrichment in the universal DHS regions combined by ENCODE project.

### Transcription factor binding site analysis

The core xenobiotic response element (XRE) sequence recognized by the AHR is 5’-T/GCGTG-3’. Position specific weight matrices (PWM) of XRE were extracted from the database TRANSFAC 7.0. The Transcription Element Search System (TESS) was used with default setting to search for AHR binding sites in the promoter region of each inhibitory gene (53).

### Chromatin immunoprecipitation assay

Chromatin immunoprecipitation assay was performed on CD8^+^ T cells activated with anti-CD3/anti-CD28 in the presence of kynurenine (50μM) or TCDD (10nM), using the EpiTect ChIP kit (Qiagen) according to the manufacturer’s protocol. Cells were crosslinked in 1% formaldehyde at 37°C for 10 minutes. After chromatin shearing by sonication, the sheared chromatin was precleared with Protein A beads, and then incubated overnight with a purified anti-AHR antibody (BioLegend) while rotating at 4°C. After DNA isolation and purification, the IP-DNA was quantified by real-time PCR. The following primer sequences were used for analysis of AHR binding to DNA: PDCD1 forward 5’-TATTTGAGGAAGGCATGAGC-3’, reverse, 5’-TCTTAACACACACGCAATCC-3’.

### Statistics

Flow cytometry, PCR, and confocal microscopy results are expressed as mean ± SEM and analyzed by two-tailed, Student’s t-test, or one- and two-way ANOVA. Kaplan-Meier curves for survival analysis were analyzed by the Log-rank Test. Statistical significance was determined by a p<0.05. GraphPad Prism 7 software was used to perform analyses. The Metabolomics data was log2-transformed and normalized with internal standard per-sample, per-method basis. For every metabolite in the normalized dataset, two-sample t-tests were conducted to compare expression levels between different groups. Differential metabolites were identified by adjusting the p-values for multiple testing at an FDR<0.25 and generated a heat map. The transformed and normalized levels of metabolites were visualized by heat map. Metabolites were grouped by hierarchical clustering using Euclidean distances and complete linkage. Comparison of group means was performed by independent sample t-tests and two-way ANOVA. Metabolic pathways were obtained from Kyoto Encyclopedia of Genes and Genomes (KEGG) data base (54). The overall difference of groups of metabolites between conditions was analyzed by GlobalANCOVA (55). Principal component analysis (PCA) was performed in standard approach. Variables and subjects were projected onto the plane of first two components to generate scatter plots. The differential gene expression analysis was performed by using linear models and empirical Bayesian methods as implemented in R limma package (56). Hypergeometric tests were used for pathway enrichment analysis as implemented in R ReactomePA package (57). Blood transcriptome modules (BTMs) were obtained from the work by Li *et al* (35). Similar to the analysis of metabolism, the overall difference of genes in each module between conditions was analyzed by GlobalANCOVA. For heat map visualization and PCA of BTM levels, the expression of all genes in module was averaged to form an overall expression level for that module. Clustering analysis of module activities were performed in same approach as that for metabolic data. The Luminex measurement of cytokines and chemokines were log-transformed before analysis. For each mice strain, the mean levels of each group were compared by two-way ANOVA, with treatment group and time as the two factors. All tests were two-sided. False discovery rates (FDRs) for transcriptomic analysis were calculated using Benjamini-Hochberg method (58).

### Study approval

All animal experiments were carried out according to protocol guidelines reviewed and approved by the Institute Animal Care and Use Committee (IACUC) of Roswell Park Comprehensive Cancer Center (Buffalo, NY).

## Author contributions

AAM, RM, TT, FQ, RYH, AAL and KO designed the experiments. AAM and KO wrote the manuscript. AAM performed the majority of the experiments and analyzed the data. RM and TT performed some of the experiments. VP and NP performed the metabolomics assay. PKS performed genomic assays. AAM, RM, HY, TL, JW, SB, and SL performed computational, bioinformatics, and statistical analyses. All authors discussed the results and commented on the manuscript.

## Acknowledgements

This work was supported by the National Cancer Institute (NCI) funded RPCI-UPCI Ovarian Cancer SPORE P50CA159981, NIH U24 CA232979, NIH R01 CA158318, NCI Cancer Center Support Grant P30 CA016056, NCI P30CA016056, NIH/NCI R01CA220297, and NIH/NCI R01CA216426 grants, and the use of Roswell Park Comprehensive Cancer Center’s shared resources including the Flow Cytometry Core, Laboratory Animal Resource, Biostatistics Shared Resource, Bioinformatics Shared Resource, and Genomics Shared Resource. We would like to thank Cheryl Eppolito for breeding animals used in this study, and the James N. Jarvis MD Lab at the University at Buffalo Jacobs School of Medicine and Biomedical Sciences for providing their expertise with the ATAC-seq experiment.

