## Supplemental tables and figures for "Indoleamine 2,3-dioxygenase upregulates PD-1 expression on ovarian tumor infiltrating CD8^+^ T cells via kynurenine activation of the aryl hydrocarbon receptor"

### Supplemental Figure 1

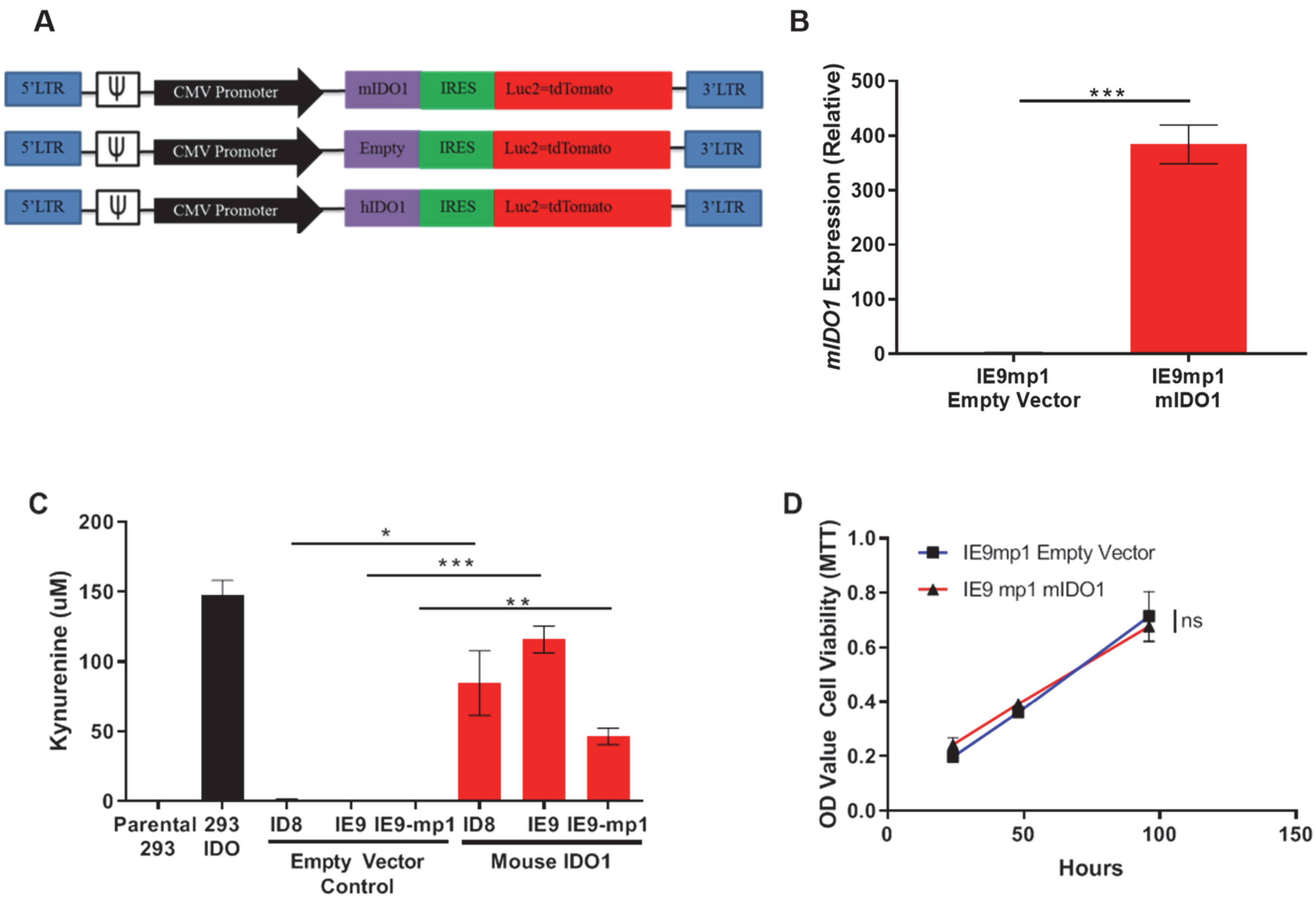


<u>Supplemental Figure 1:</u> **Established a mouse epithelial ovarian cancer cell line which expresses IDO1 and demonstrates IDO1 functional enzyme activity. (A)** Schematic representation of retroviral vector constructs encoding full length mouse IDO1 cDNA transfected into parental ID8, IE9 or IE9mp1 tumor cell lines. Luc2-dTomato, a marker for the mIDO1 gene and empty-control vectors. **(B)** mIDO1 gene expression in IE9mp1-mIDO1 and IE9mp1-EV control tumor cells by q-PCR. Results presented relative to the expression of *GAPDH*. **(C)** Kynurenine concentrations measured in mIDO1 and empty vector control ID8, IE9 and IE9-mIDO1 tumor cells, and parental HEK-293 and HEK-293-IDO cell culture supernatants by colorimetric assay. **(D)** in vitro cell viability of IE9mp1-mIDO1 or IE9mp1-EV tumor cells was measured by MTT assay at 24, 48 and 96 hrs. O.D. is Optical Density. $*p < 0.05$, $**p < 0.01$, $***p < 0.001$, ns: not significant, by Student's t test **(B, C** and **D)**. The data represent means ± SEM of three independent experiments performed in triplicate.

### Supplemental Figure 2

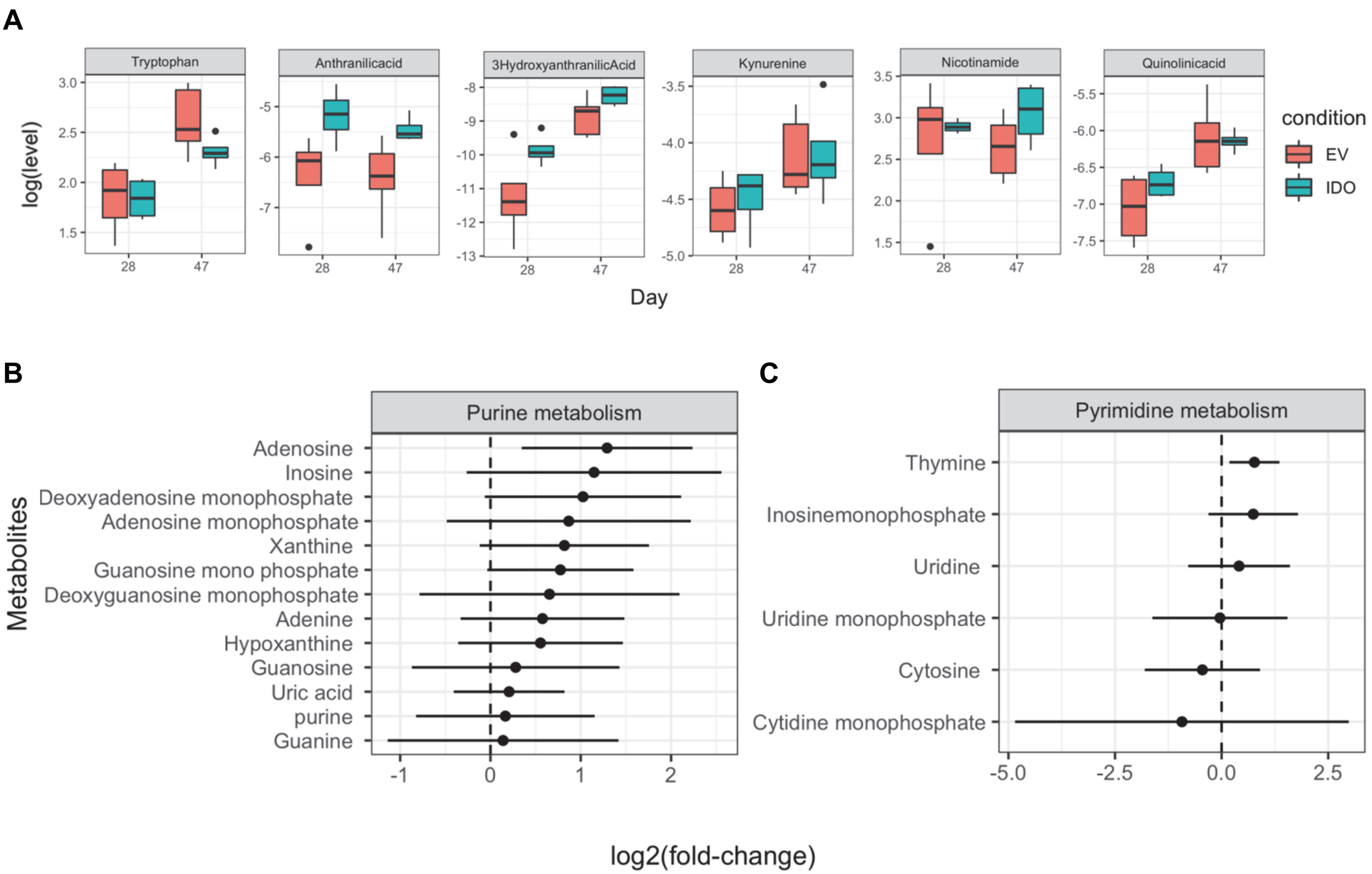


**Supplemental Figure 2: Metabolites levels from the kynurenine, nicotinamide, purine and pyrimidine metabolic pathways are affected by IDO1 expression by the tumor. (A)** Boxplot of tryptophan, anthranilic acid, 3-hyrdroxyanthranilic acid, kynurenine, nicotinamide, and quinolinic acid metabolite levels measured in IE9mp1-mIDO1 and IE9mp1-EV tumor from WT C57BL/6 mice at Days 28 and 47. **(B)** 95% confidence interval of the mean log-fold change in metabolites of the purine **(C)** and pyrimidine metabolism pathways in IE9mp1-mIDO1 and IE9mp1-EV tumor from WT C57BL/6 mice at Days 28 and 47.

#### Supplemental Figure 3

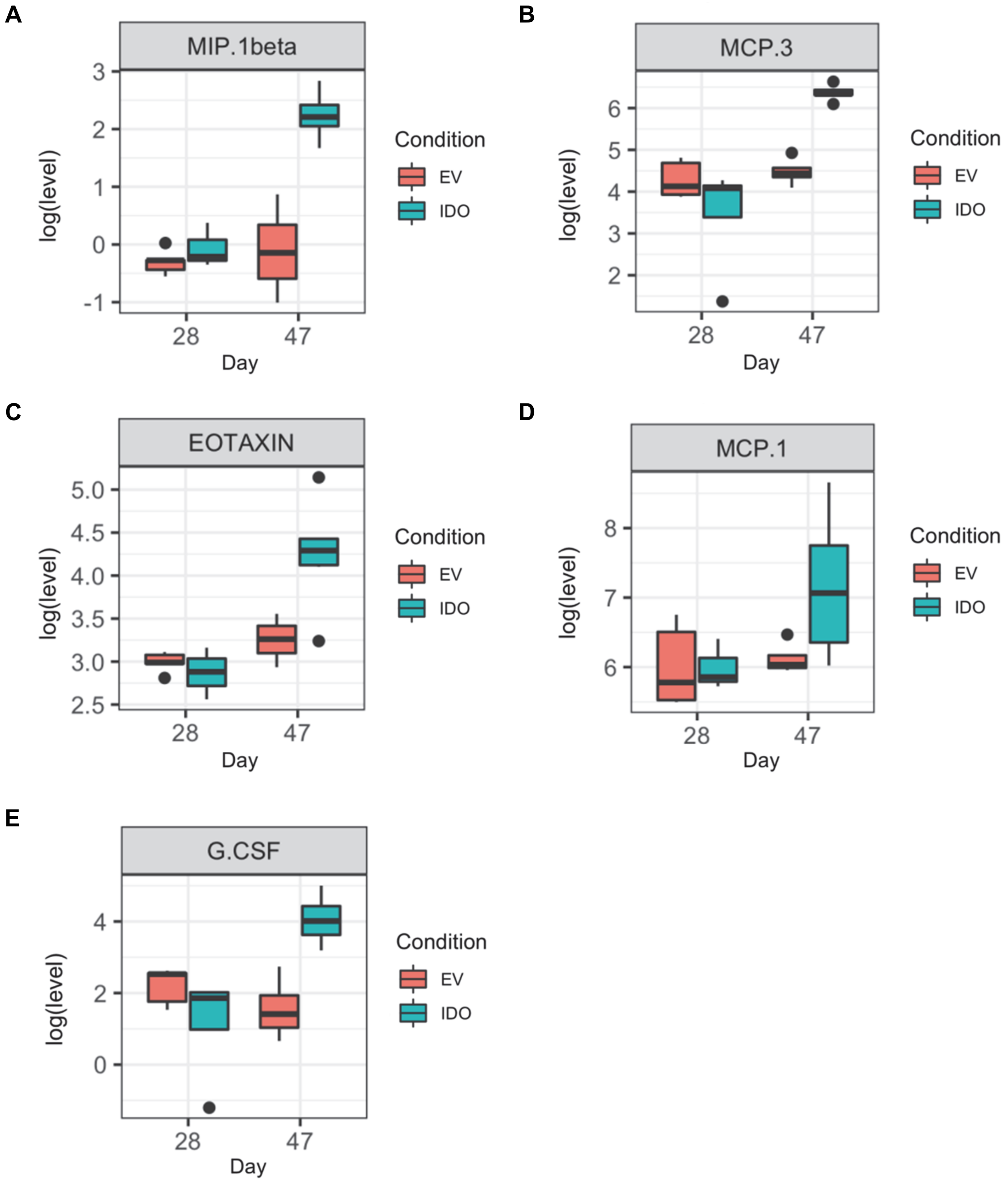


**<u>Supplemental Figure 3:</u> Tumor-derived IDO1 mediates upregulation of suppressive chemokines in WT C57BL/6 mice tumor ascites, compared to tumors which lack IDO1. (A)** MIP1-beta (CCL4) *P*=0.0058, **(B)** MCP-3 (CCL7) *P*=0.00047, **(C)** Eotaxin (CCL11) *P*=0.0094, **(D)** MCP-1(CCL2) *P*=0.055, and **(E)** G-CSF *P* =0.0044 were measured in cell-free tumor ascites fluid from IE9mp1-EV and IE9mp1-mIDO1 tumor-bearing WT C57BL/6 mice at Days 28 and 47 by ProcartaPlex Luminex immunoassay. Statistical significance at Day 47 was determined by t-tests following two-way ANOVA **(A, B, C, D** and **E)**.

### Supplemental Figure 4

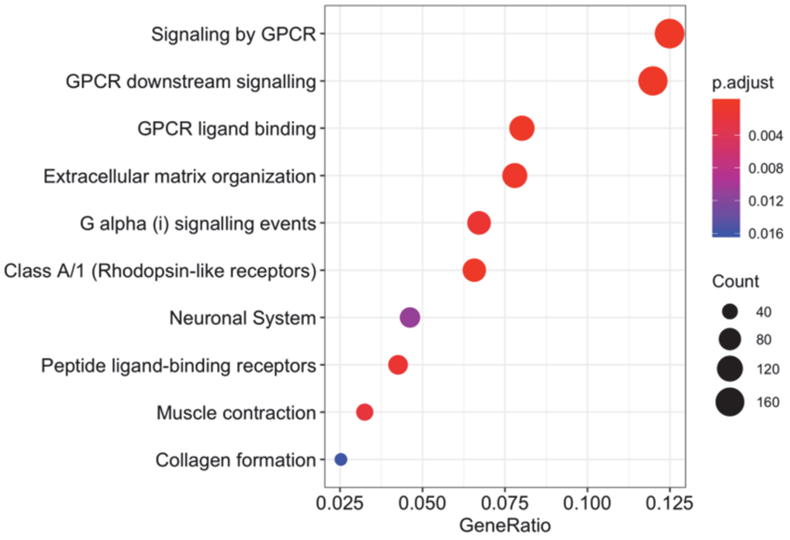


**Supplemental Figure 4: Tumoral IDO1 mediates transcriptomic changes tumor infiltrating lymphocytes.** Pathway Enrichment Analysis in CD8$^{+}$ T cells of IE9mp1-mIDO1 tumors revealed 244 modules enriched in GPCR downstream signaling in IDOKO mice at Day 48. Gene Ratio is the number of genes enriched divided by the total number of genes in each pathway. Circles represent the number of genes. P-values are showed in color scale.

### Supplemental Figure 5

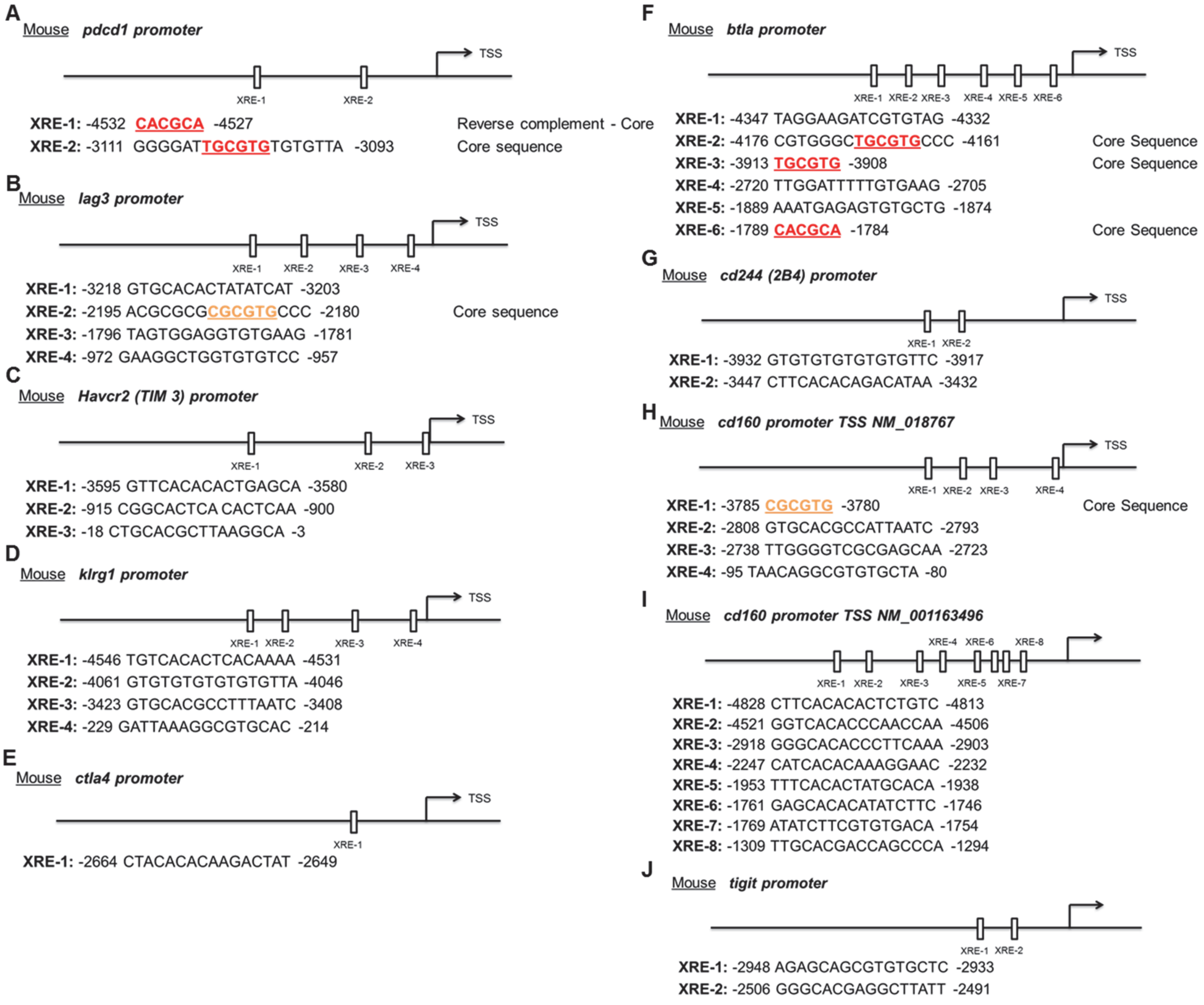


**Supplemental Figure 5: Mouse inhibitory receptor genes contain aryl hydrocarbon receptor (AHR) binding sites. (A)** AHR responsive element binding sites in the promoter region of mouse *pdcd1*, **(B)** *lag3*, **(C)** *Havcr2*, **(D)** *klrg1*, **(E)** *ctla4*, **(F)** *btla*, **(G)** cd244, **(H** and **I)** *cd160*, and **(J)** *tigit* genes. TGCGTG and CACGCA (reverse complement) shown in red, or CGCGTG and CACGCG (reverse complement) shown in orange.

### Supplemental Figure 6

**A**

Human ***pdcd1 promoter***

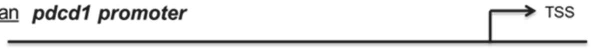


**XRE-1:** -3691 AAGGGTGGGTGTGCCA -3676
**XRE-2:** -2732 TGGAAATCGTGTGTTG -2717
**XRE-3:** -2718 GGGGTGTGGTGTGTTG -2703
**XRE-4:** -2687 GACTGTGTGTGTGTTG -2672
**XRE-5:** -2669 GATTGTGTGTGTGTTG -2654
**XRE-6:** -2620 TTTTGTGTGTGTGTTG -2605
**XRE-7:** -2566 TTTTGTGTGTGTGTTG -2551
**XRE-8:** -2513 TGAGGATTGTGTGCAG -2498
**XRE-9:** -2499 TTGTGTGTGTGTGTTG -2484
**XRE-10:** -2463 GATTGTGTGTGTGTTG -2448
**XRE-11:** -2446 AGTTGTGTGTGTGTTG -2431
**XRE-12:** -2394 TTTTGTGTGTGTGTTG -2379
**XRE-13:** -2340 AATTGTGTGTGTGTTG -2325
**XRE-14:** -2273 TTTTGTGTGTGTGTTG -2258
**XRE-15:** -2233 TGTGGATTGCGTGTAC -2218 Core Sequence
**XRE-16:** -2179 AGAGGATTGTGTGTTG -2164
**XRE-17:** -2113 ATTTGTGTGTGTGTTG -2098
**XRE-18:** -2078 AATTGTGTGTGTGTTG -2063
**XRE-19:** -2026 TTCTGTGTGTGTGCTG -2011
**XRE-20:** -1990 GATTGTGTGTGTGTTG -1975
**XRE-21:** -1956 TTGTGTGTGTGTGCTG -1941
**XRE-22:** -1954 GGAGATTGTGTGTGTGTGC -1936
**XRE-23:** -1920 GGTTGTGTGTGTGTTG -1905
**XRE-24:** -1872 TGTAGATTGCGTGTTG -1857 Core Sequence
**XRE-25:** -1835 TTTTGTATGTGTGTTG -1820
**XRE-26:** -1797 TGGAAATCGTGTGTTG -1782
**XRE-27:** -1783 GGAGTTGTGTGTGTTG -1768
**XRE-28:** -1750 GAAAGTGTGTGTGCTG -1735
**XRE-29:** -1264 GTGCACGCCTGTGGTC -1249
**XRE-30:** -750 GGGAGGTGGGTGTATTGCC -732
**XRE-31:** -622 CACGCG -617 Core Sequence

**B**

Human ***lag3 promoter***

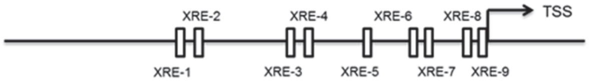


**XRE-1:** -4336 CTTCACGCTCCCTAAC -4321
**XRE-2:** -4303 TGT**CACGCA**CACAGGT -4288 Core Sequence
**XRE-3:** -3952 CTGGGTCTCGTGTCATCCC -3934
**XRE-4:** -3576 CAGCACACTCCTACCA -3561
**XRE-5:** -2010 GTTCACACACCTCTAA -1995
**XRE-6:** -1109 GATCACACAAGGACAC -1094
**XRE-7:** -1088 TGGCACACCAACACCA -1073
**XRE-8:** -949 AGGGAATTGAGTGCCC -934
**XRE-9:** -474 **CGCGTG** -469 Core Sequence

**C**

Human ***Havcr2 (TIM 3) promoter***

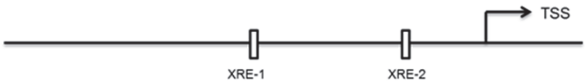


**XRE-1:** -3270 GGAGAATTGCTTGAAC -3255
**XRE-2:** -1760 TTGCACACTAAAGTAC -1745

**D**

Human ***klrg1 promoter***

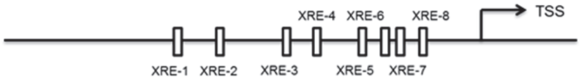


**XRE-1:** -4549 GCATATGCACCCAACAGTC -4531
**XRE-2:** -3288 TATTAAGGGCGTGAAA -3273
**XRE-3:** -2761 GTGCACACACTGAGGA -2746
**XRE-4:** -2190 GAT**CACGCG**GATGGTT -2175 Core Sequence
**XRE-5:** -1606 GTT**CACGCA**GTGAACC -1591 Core Sequence
**XRE-6:** -750 GAAGACTTGTGTGACC -735
**XRE-7:** -566 TTTTTTTTTCGTGAAC -551

**E**

Human ***ctla4 promoter***

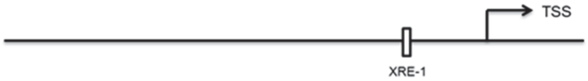


**XRE-1:** -2502 GGTGGGTGCCTGCAATCCC -2484

**F**

Human ***btla promoter***

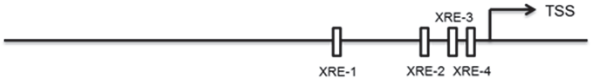


**XRE-1:** -1638 CTTCACTCAACAATCA -1623
**XRE-2:** -403 **TGCGTG** -398 Core Sequence
**XRE-3:** -320 CTGCACACTAGTCTCT -305
**XRE-4:** -299 GTGCACACCCTAATAA -284

**G**

Human ***cd244 (2B4) promoter***

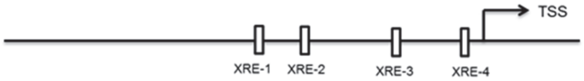


**XRE-1:** -4441 GTGCACGCCACCCAGC -4426
**XRE-2:** -4349 GTGCAGTGGCGTGATC - 4334
**XRE-3:** -2297 TTGGGTCCTCGTGCTA -2282
**XRE-4:** -130 TATCACGAGACTAGCA -115

**H**

Human ***cd160 promoter***

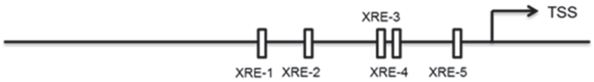


**XRE-1:** -4173 GATCACGCCACTGTAC -4158
**XRE-2:** -3158 GGTCACACCATTCTCC -3143
**XRE-3:** -2423 TGGCACACAATAACCT -2408
**XRE-4:** -2422 GGCCTGGCACACAATAACC -2404
**XRE-5:** -1403 TGAGACTTGCCTGCCA -1388

**I**

Human ***tigit promoter***

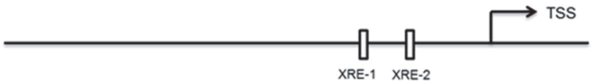


**XRE-1:** -2104 GAGCACGACCCCCTGC -2089
**XRE-2:** -1778 **CGCGTG** -1773 Core Sequence

**Supplemental Figure 6: Human inhibitory receptor genes contain aryl hydrocarbon receptor (AHR) binding sites. (A)** AHR responsive element binding sites in the promoter region of mouse *pdcd1*, **(B)** *lag3*, **(C)** *Havcr2*, **(D)** *klrg1*, (E) *ctla4*, **(F)** *btla*, **(G)** cd244, **(H)** *cd160*, and **(I)** *tigit* genes. TGCGTG and CACGCA (reverse complement) shown in red, or CGCGTG and CACGCG (reverse complement) shown in orange.

### Supplemental Figure 7

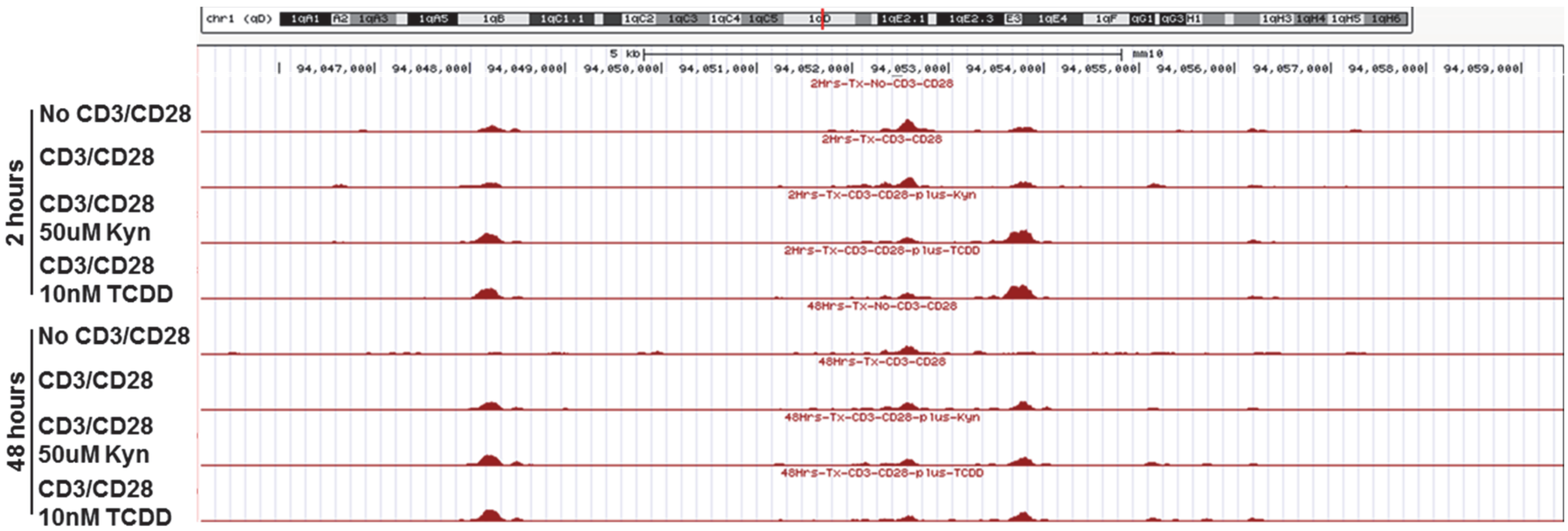


<u>**Supplemental Figure 7:**</u> **ATAC-seq identifies genome-wide chromatin accessibility in regulatory regions of the LAG3 gene in kynurenine-treated CD8⁺ T cells.** UCSC Genome Browser plot of chromatin accessible regions in DNA regulatory elements on the LAG3 gene by ATAC-seq analysis from WT C57BL/6 mice CD8⁺ T cells activated by anti-CD3/CD28 and treated with KYN (50μM) or TCDD (10nM) for 2 and 48 hrs.

**Supplemental Figure 8**

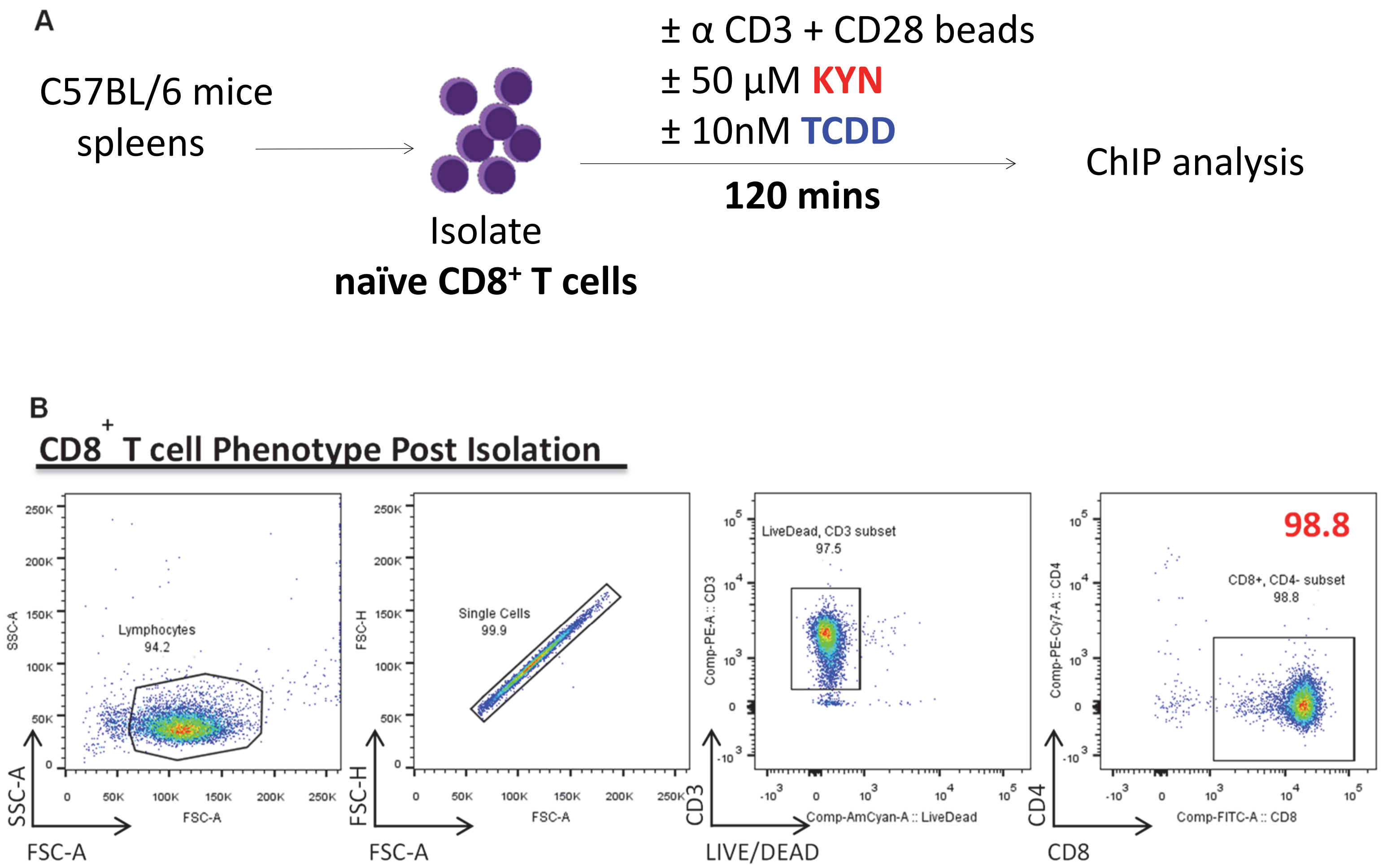


•**Supplemental Figure 8: CD8+ T cells isolated by negative selection from WT C57BL/6 mice spleens. (A)** Experimental design used in the chromatin immunoprecipitation (ChIP) assay. Naïve CD8+ T cells from WT C57BL/6 spleens were sorted by magnetic bead negative-selection, anti-CD3/CD28 bead-activated and treated with KYN (50μM) or TCDD (10nM) for 2 hrs. **(B)** High purity of CD3+CD8+ T cells were confirmed by flow cytometry before preforming the CHIP assay. Numbers in outlined areas indicate percentage of gated populations.

#### Supplemental Figure 9

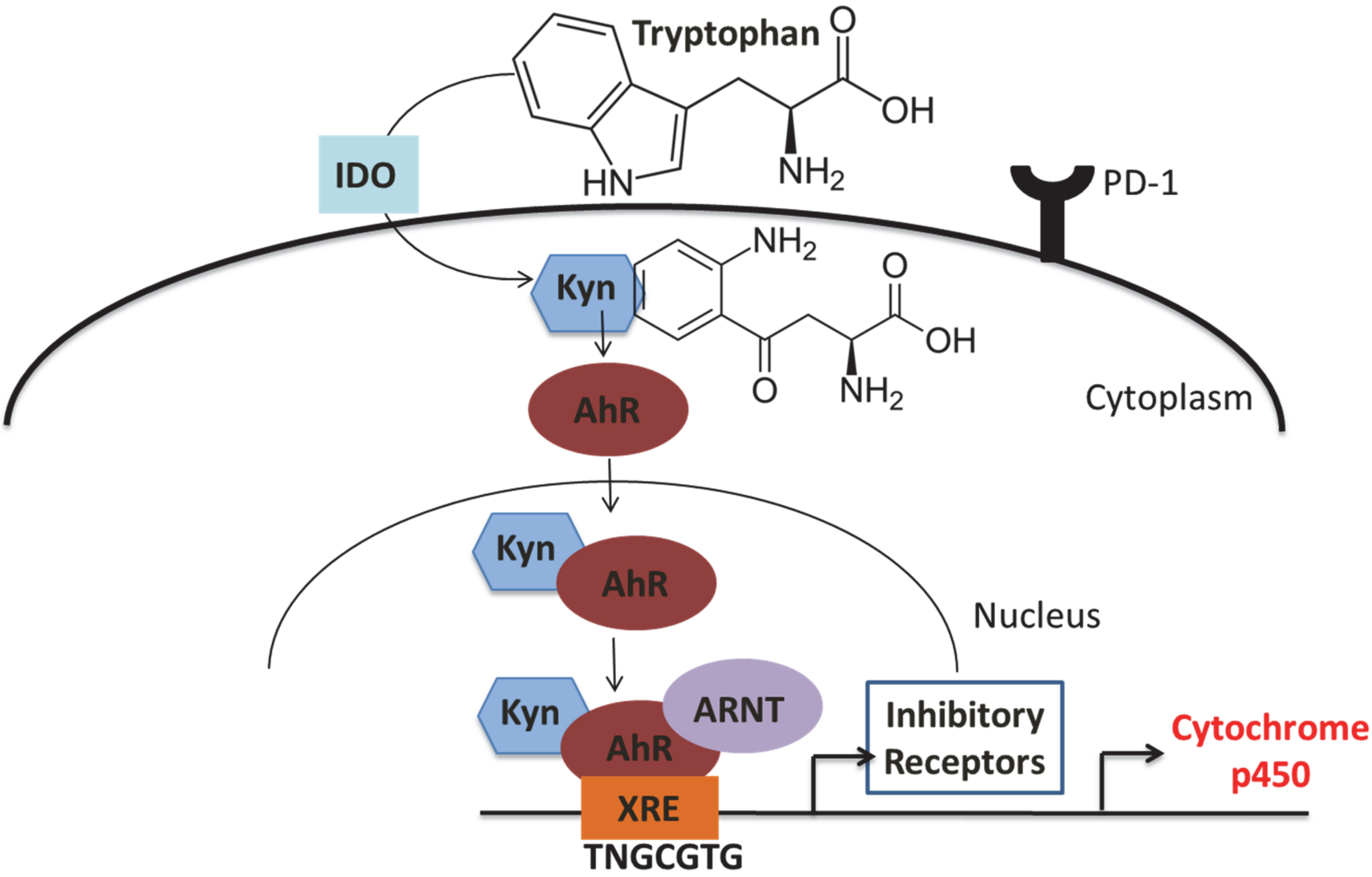


**Supplemental Figure 9:** *Model: Kynurenine mediated aryl hydrocarbon receptor activation will contribute to induction PD-1 expression on CD8⁺ T cells.* **(A)** Following catabolism of tryptophan by tumor-derived IDO, the metabolite kynurenine was generated and thus, ligated and activated the transcription factor AHR to translocate to the nucleus and bind to XRE sequences in the PD-1 gene promoter. Kynurenine induced PD-1 expression on $CD8^+$ T cells in an AHR-dependent manner demonstrated by expression of the downstream AHR target gene, CYP1a1.

### Supplemental Table 1

| | |
|---|---|
| AHR | EHHADH |
| AANAT | EIF2AK4 |
| ACAT1 | GCDH |
| ACAT2 | GZMB |
| ALDH1B1 | HAAO |
| ALDH2 | HADH |
| ALDH3A2 | HADHA |
| ALDH7A1 | IDO1 |
| ALDH9A1 | IL2 |
| AHR | IDO2 |
| AOC1 | KMO |
| AOX1 | KYNU |
| ASMT | MAOA |
| CAT | MAOB |
| CCBL1 | OGDH |
| CCBL2 | OGDHL |
| CD3E | TDO2 |
| CD8A | TPH1 |
| CYP1A1 | WARS |
| CYP1A2 | WARS2 |
| CYP1B1 | 3HAO |
| DDC | ACMSD |
| ECHS1 | QPRT |

•**Supplemental Table 1: Genes related to the tryptophan metabolism pathway and the AHR signaling pathway.** List of 44 genes related to the tryptophan metabolism pathway and the AHR signaling pathway, and tumor infiltrating lymphocyte genes CD3E, CD8A, Granzyme B, and IL-2 analyzed in the TCGA dataset.

### Supplemental Table 2

| ID | Module title | Size | Module category |
|---|---|---|---|
| *M4.8* | cell division - E2F transcription network | 19 | biological process |
| *M5.0* | regulation of antigen presentation and immune response | 81 | immune |
| *M5.1* | T cell activation and signaling | 25 | immune |
| *M7.0* | enriched in T cells (I) | 62 | immune |
| *M7.1* | T cell activation (I) | 50 | immune |
| *M7.2* | enriched in NK cells (I) | 49 | immune |
| *M7.3* | T cell activation (II) | 31 | immune |
| *M7.4* | T cell activation (III) | 14 | immune |
| *M10.1* | E2F1 targets (Q4) | 21 | TF targets |
| *M12* | CD28 costimulation | 10 | immune |

**Supplemental Table 2: The first 10 out of 103 modules identified in Cluster 1 from the transcriptomics analysis of TILs.** List revealed genes related to regulation of antigen presentation and immune response, T cell activation and signaling, and CD28 co-stimulation.

### Supplemental Table 3

| Gene | KO | | WT | |
|---|---|---|---|---|
| | Log FC | p-value | Log FC | p-value |
| *BTLA* | -4.3419 | 0.0013 | 1.2695 | 0.1906 |
| *CD160* | -3.7265 | 0.0299 | 0.0612 | 0.9643 |
| *CTLA4* | -1.7608 | 0.2041 | 2.2533 | 0.0795 |
| *KLRG1* | 3.0118 | 0.0497 | -1.9533 | 0.1383 |
| *LAG3* | 3.1198 | 0.0070 | -0.1990 | 0.8181 |
| *PDCD1* | 4.4151 | 0.0106 | -1.9362 | 0.1606 |
| *TIGIT* | 2.3714 | 0.0600 | -0.9822 | 0.3528 |

**Supplemental Table 3: Tumoral IDO1 led to upregulation of PDCD1 inhibitory receptor gene expression on CD8$^{+}$ TILs.** Data shown are log fold-change and p-values for inhibitory receptor gene expression that are significantly decreased (blue) and increased (red) in IE9mp1-mIDO1 tumor-bearing WT C57BL/6 and IDOKO mice.
